# Substrate binding modulates the conformational kinetics of the secondary multidrug transporter LmrP

**DOI:** 10.1101/2020.04.09.034439

**Authors:** Aurélie Roth, Chloé Martens, Thomas van Oene, Anders Barth, Simon Wanninger, Don C. Lamb, Jelle Hendrix, Cédric Govaerts

## Abstract

The Major Facilitator Superfamily (MFS) is the largest family of secondary active membrane transporters and is found in all domains of Life. MFS proteins are known to adopt different conformational states, yet details on the interconversion rates are crucially needed to understand or target their transport mechanism. Here, we studied the proton/multidrug antiporter LmrP as a model system for antibiotic resistance development in bacteria. The conformational cycle of LmrP is triggered by the protonation of a network of specific amino acids, yet the role of the transported substrate in these transitions has been puzzling. To measure LmrP structure in real-time, we performed solution-based single-molecule Förster resonance energy transfer (smFRET) using a confocal microscope with direct alternating donor/acceptor excitation and multiparameter (intensity, lifetime, anisotropy) detection. Lowering pH from 8 to 5 triggered an overall conformational transition, corroborating that detergent solubilization allows studying the LmrP transport cycle using smFRET. Using a newly developed linear 3-state photon distribution analysis (PDA) model, we show that the apo protein interconverted between two structures at low rate (>>10 ms dwell time) at the cytosolic side while it interconverts dynamically between the 3 states (< 10 ms dwell time) at the extracellular side. When the Hoechst 33342 model substrate is added, inward conformational interconversions are greatly accelerated, coupled to an overall outward conformational halting, consistently with efficient proton exchange with the extracellular environment. Roxithromycin substrate binding did not halt but shift conformational interconversions from one pair of states to another. Substrate dependent structural heterogeneity is indicative of a general mechanism by which MFS transporters can efficiently transport a variety of substrates, and advocates for combined structure/dynamics-based drug design when targeting MDR transporters.

**BRIEF SUMMARY:** We studied the conformational cycle of LmrP, a model for multidrug efflux pumps, using single-molecule Förster resonance energy transfer (smFRET). By following changes in FRET signal between different sets of positions, we specifically investigated how substrate binding modulates structural conversions between inward-open and outward-open states. Using a newly developed probabilistic analysis for describing sequential interconversion kinetics, we show that the apo protein slowly interconverts between defined states at the cytosolic and at the extracellular sides. Binding of the model substrate Hoechst33342 leads to an increase in conformational interconversions at the intracellular side while the extracellular side shows a drastic decrease in conversion, indicating a kinetic uncoupling between both sides. Remarkably, binding of roxithromycin, while also increasing interconversion on the intracellular side, did not slow the extracellular conversions. This indicates that multidrug pumps have evolved substrate-dependent transport mechanisms than enable transport of structurally diverse collection of substrates.

## I. Introduction

Membrane transporters allow the passage of different molecules across the cell membrane by undergoing large conformational changes to alternatively expose the substrate to the extracellular or the cytosolic side [1]. More particularly, transporters of the Major Facilitator Superfamily (MFS) follow the rocker-switch model in which the two domains of the protein pivot around the binding site [2,3]. MFS form the largest family of membrane transporters and are present in all domains of life [3,4]. These transporters can be uniporters, symporters and antiporters, and the substrates transported are varied, e.g. sugar, drugs, amino acids, peptides, lipids, anions and cations [3,4]. In bacteria, proteins from this family are involved in the uptake of nutrients and extrusion of deleterious compounds [5]. The MFS also contains many efflux pumps that can recognize cytotoxic compounds, e.g. antibiotics, and prevent them from reaching their target in the cell [6–8]. Such multidrug efflux pumps significantly contribute to antibiotic resistance (MDR) in pathogenic bacteria [9,10].

To better understand the conformational changes of MFS transporters, we focused on LmrP from *Lactococcus lactis*. This multidrug efflux protein exports several hydrophobic cytotoxic compounds in exchange of protons in an electrogenic way [11]. Using distance electron electron resonance (DEER) studies, we have previously investigated the coupling between the conformational changes of LmrP and the passage of the protons and of the substrate, in detergent micelles [12] and in a lipid bilayer [13]. The transporter transitions from an outward-open to an inward-open state depending on the pH, and specifically on the protonation of networks of specific amino acids, especially the D68 [12,13] that is part of the “motif A” highly conserved through MFS transporters [14]. The prototypical substrate Hoechst 33342 stabilizes an outward-open state [12], which has been confirmed recently with the X-ray structure of the protein in complex with this substrate in the outward-open conformation (Debruycker et al, under revision)[ref structure LmrP]. These measurements offered a snapshot of all the conformations adopted by the protein at a given time [15] but likely miss transitions, which are by definition transient states, that are key to understand the transport cycle at a molecular level. To adequately describe transport cycles it is, however, important to characterize both the structural but also kinetic aspects of possible transitions between conformers. Indeed, key structural states may be very shortlived and therefore represent only a small subset of the conformational ensemble at equilibrium. Finally, although simulation techniques can provide a molecular description of short-lived states, a fully experimental kinetic characterization of transition states is highly desirable.

Single-molecule Förster Resonance Energy Transfer (smFRET) allows to characterize intramolecular distances between a donor (D) and an acceptor (A) fluorophores attached on individual proteins on timescales ranging from ns to minutes. In particular, fast time scales can be followed using Pulsed Interleaved Excitation (PIE) with which the sample in solution is analyzed on a confocal microscope (Fig. 1B) and excited with two lasers (corresponding to the D or A fluorophore) that alternate on the ns timescale. The combination of multiparameter fluorescence detection (MFD) and PIE allows the simultaneous monitoring of the key fluorescent parameters (the quantum yield, the fluorescence lifetime, the fluorescence intensities and the fluorescence anisotropy) that describe the local environment of the fluorophores [16–18]. Most importantly the efficiency of the energy transfer (*E*) between the donor and the acceptor is retrieved, which is inversely proportional to the distance between the probes and thereby reflects the conformations adopted by the protein (Fig. 1D) [19–21]. The single molecule detection allows the identification of dynamic molecules as well as conformational heterogeneity and transient states that are usually lost in ensemble measurements techniques such as FRET or DEER [22]. Therefore, smFRET allows monitoring the dynamics and kinetics of protein motion. Using Photon Distribution Analysis (PDA) we can quantify the interconversion rate of the protein on the millisecond timescale [25,26].

**Fig.1:**
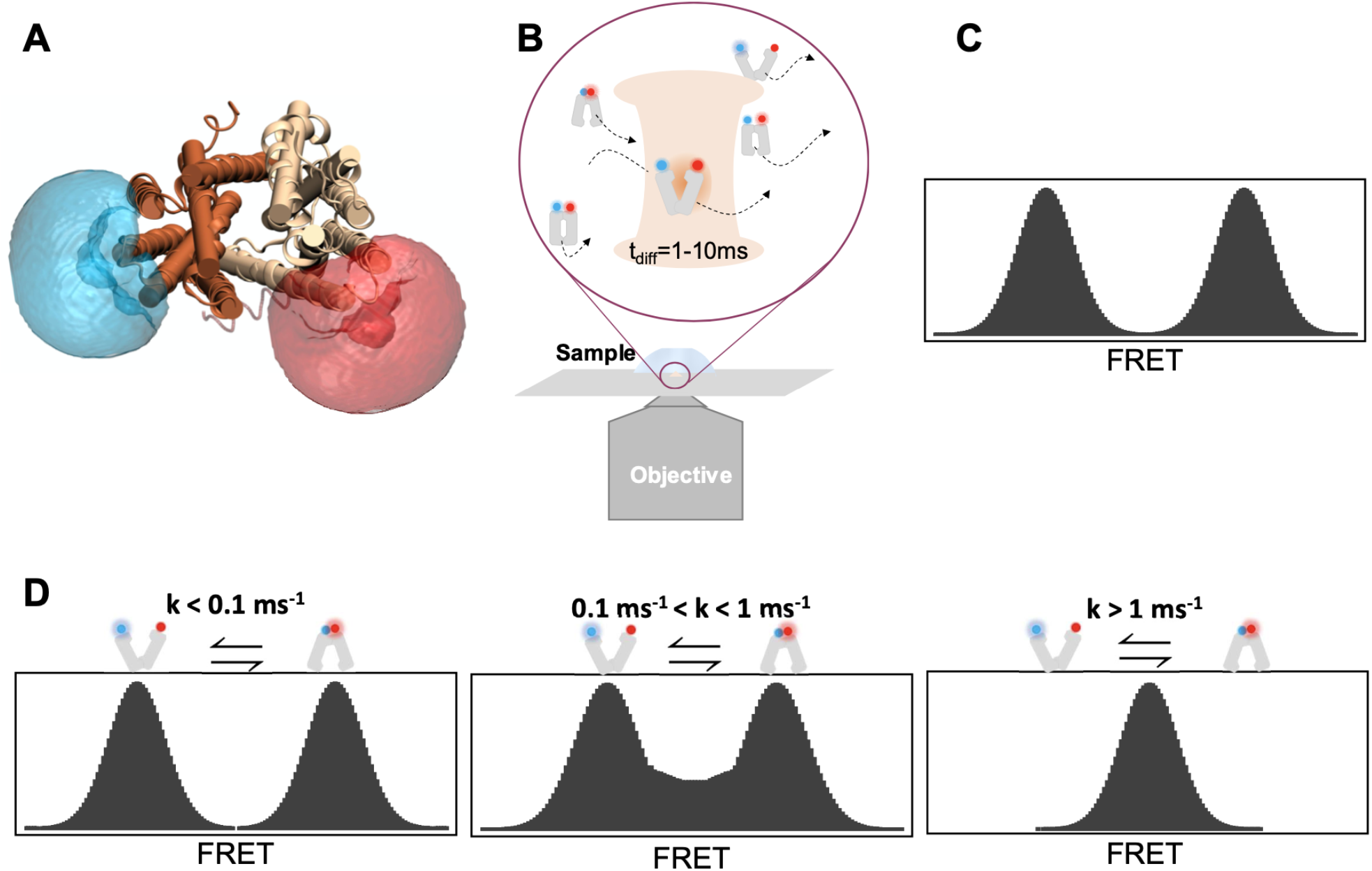
Principles of the smFRET measurements. **A:** Cartoon representation of LmrP 98C (TM3)-374C (TM11) labelled with ATTO488 (blue) and Alexa647 (red) respectively. The N- and C-halves, composed of TM1-6 and TM7-12 respectively, are represented in dark and light brown respectively. The probe clouds were calculated using the FPS software [33]. All LmrP representations were prepared using VMD [34]. **B:** Illustration of proteins interconverting between an outward-open/inward-closed and outward-closed/inward-open conformation and diffusing through a confocal volume during 1-10ms. **C:** Example of a burst measurement obtained with a confocal microscope. The donor emission after excitation of the donor is represented in green. The FRET bursts after the donor excitation are represented in red. **D:** The *E* histograms retrieved depend on the different conformations adopted by the protein but also on the interconversion rate. In the first case, if the protein is not interconverting when diffusing into the confocal volume (k< 0.1ms^−1^), the two populations corresponding to the high and low FRET states are clearly separated. If the proteins are interconverting on the timescale of their diffusion (0.1 ms^−1^ < k < 1ms^−1^), the two FRET populations are connected by a smear corresponding to the molecules interconverting during their diffusion into the focus. Finally, if the proteins are interconverting between the two conformations faster than their diffusion through the focus, only the interconverting molecules are detected resulting in an average FRET population.

In this study, we used smFRET/MFD-PIE to characterize the conformational dynamics of LmrP on the ms timescale with a newly developed three-state PDA model. We observed that the conformational equilibrium of LmrP can be modulated by pH, protonation-mimetic mutations and with the addition of Hoechst. Specifically, we investigated the role of ligands on the transition kinetics between the different conformations adopted by the protein.

## II. Results

### smFRET reports on the conformational landscape and substrate binding of LmrP

To follow LmrP conformational changes in real-time with smFRET, pairs of residues were replaced by cysteines for a specific reaction with the reactive maleimide moieties of the ATTO 488 and Alexa Fluor 647 fluorescent dyes. The residues were selected to be at the end of transmembrane helices, either on the cytosolic or the extracellular side, and predicted to provide maximal conformational freedom for the attached dyes. Two cytosolic (9C-404C, 188C-404C) and three extracellular (98C-374C, 164C-312C, 164C-374C) distance reporters were selected to probe distance fluctuations across different opposing sides of the transporter between the N- and C-terminal halves (Fig. 1A). The expected FRET efficiency *E*_expected_ and corresponding distances between the probes are given in Table 1. All double cysteine mutants were functionally tested by following LmrP-mediated transport of Hoechst in *L. lactis* inverted membrane vesicles, and all exhibited a wildtype-like transport of the substrate (Fig. S1A-E). Control single-molecule experiments corroborated that changes in *E* are indeed solely related to protein conformational changes, and not to any photophysical artifact (Fig. S2).

**Table 1:** expected FRET efficiencies and corresponding distances of the closed and open conformations.

| Residues | Localisation | Outward-open<br>Inward-closed |  | Outward-closed<br>Inward-open |  |
| --- | --- | --- | --- | --- | --- |
| | | $E_{\text{expected}}$ | R (Å) | $E_{\text{expected}}$ | R (Å) |
| Cytosolic distance reporters |  |  |  |  |  |
| 9C-404C | TM1 – TM12 | 0.68 | 50.4 | 0.35 | 63.0 |
| 188C-404C | TM6 – TM12 | 0.70 | 49.5 | 0.59* | 53.4* |
| Extracellular distance reporters |  |  |  |  |  |
| 98C-374C | TM3-TM11 | 0.47 | 58.1 | 0.78 | 45.9 |
| 164C-312C | TM6 – TM10 | 0.62 | 52.4 | 0.85 | 42.4 |
| 164C-374C | TM6 – TM11 | 0.48 | 57.5 | 0.66 | 51.0 |

\* The residue 188 was absent in the inward-open model. The distance and $E_{\text{expected}}$ were calculated based on the 187C-404C mutant.
The $E_{\text{expected}}$ and the derived inter-probes distances in the outward-open/inward-closed state were calculated based on the crystal structure of LmrP binding Hoechst and the outward-closed/inward-open data were calculated based on a model of LmrP in this conformation. The position of the gamma atom, if available, of the non-mutated residue was used for the calculations.

We first tested the effect of pH on the conformational landscape of LmrP as reported by our smFRET distance reporters. At the cytosolic side of the protein at pH 8, two FRET states were observed (Fig. 2A-B, black). This was particularly obvious for the 188C-404C pair where a major state at *E* ≈ 0.6 and a minor state at *E* ≈ 0.2 were present. At the extracellular side at pH 8, one major broad state was observed, (Fig. 2C-E, black). At pH 5, the FRET signature at both sides of the protein changed dramatically. Only one main population remained at the cytosolic side (Fig. 2A-B, orange), indicating that this part of the protein opens up. Conversely, a high FRET state became apparent at pH 5 at the extracellular side (Fig. 2C-E, orange), indicating the protein had partially switched to a more closed structure. Based on the X-ray crystallography structure of LmrP in the outward-open/inward-closed state and dyes’ accessible volume (AV) calculations, we calculated *E*_expected_ for the inward-closed state to be around 0.6-0.7 for both cytosolic reporters (Table 1). Since this is in good agreement with the main FRET state at pH 8 (Fig. 2A-B, black), we conclude this state is indeed the inward-closed conformation. Similarly, on the extracellular side, the expected *E* values for the outward-open state (Table 1) were close to the main FRET state observed at pH 8 (Fig. 2C-E, black), indicating that it most likely corresponds to the outward-open state. This assignment was further confirmed by using the D68N mutation of LmrP, that is known to stabilize the outward-closed state [12] and prevent Hoechst transport in membrane vesicles of *L. lactis* [11]. Both at pH 8 and 5, the high-FRET state was indeed more populated for the mutant, confirming that it corresponds to the outward-closed conformation (Fig. S3). Together, these experiments prove that smFRET can be used to probe the different inward/outward conformational states of LmrP.

**Fig.2.**
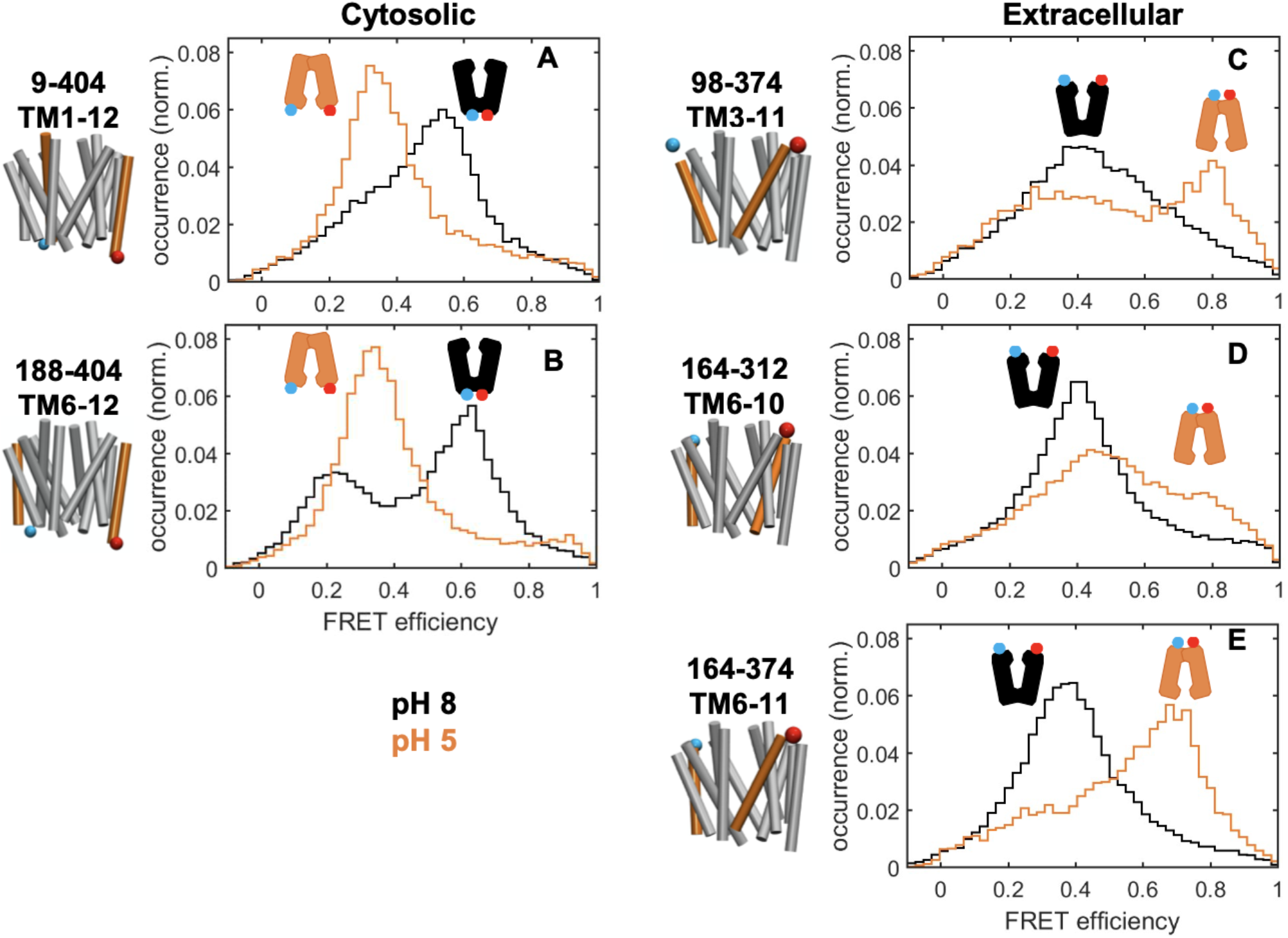
Modulation of the conformational dynamics of LmrP with the pH. Left: cartoon representations of the two constructs probing the cytosolic side and the three constructs probing the extracellular side. The helices carrying the probes (in blue (donor) and red (acceptor)) are highlighted in orange. **A-E:** For each mutant measured at pH 8 and at pH 5, the FRET histograms are shown in black and orange respectively.

Next, we tested whether the substrate Hoechst altered the conformational landscape of LmrP. In the presence of Hoechst, the cytosolic side of the protein exhibited an intermediate *E* ≈ 0.5 state at pH 8 (Fig. 3A-B). Such FRET value does not fit the 9-404 and 188-404 distances predicted for the Hoechst-bound crystal structure of LmrP (*E*_expected_ ≈ 0.68 (Table 1)). One possible explanation is that the presence of Hoechst increases the interconversion rate between different states, resulting in an ‘averaged’ FRET signature (see below and Fig. 1D). At the extracellular side Hoechst shifted the FRET histogram towards lower FRET values, suggesting the protein is able to adopt an additional ‘extra-open’ conformation (Fig. 3C-E). In summary, substrate binding to the protein was readily observed via smFRET, and suggests the substrate affects conformational dynamics of LmrP in addition to, or rather than, its mere structure.

**Fig.3.**
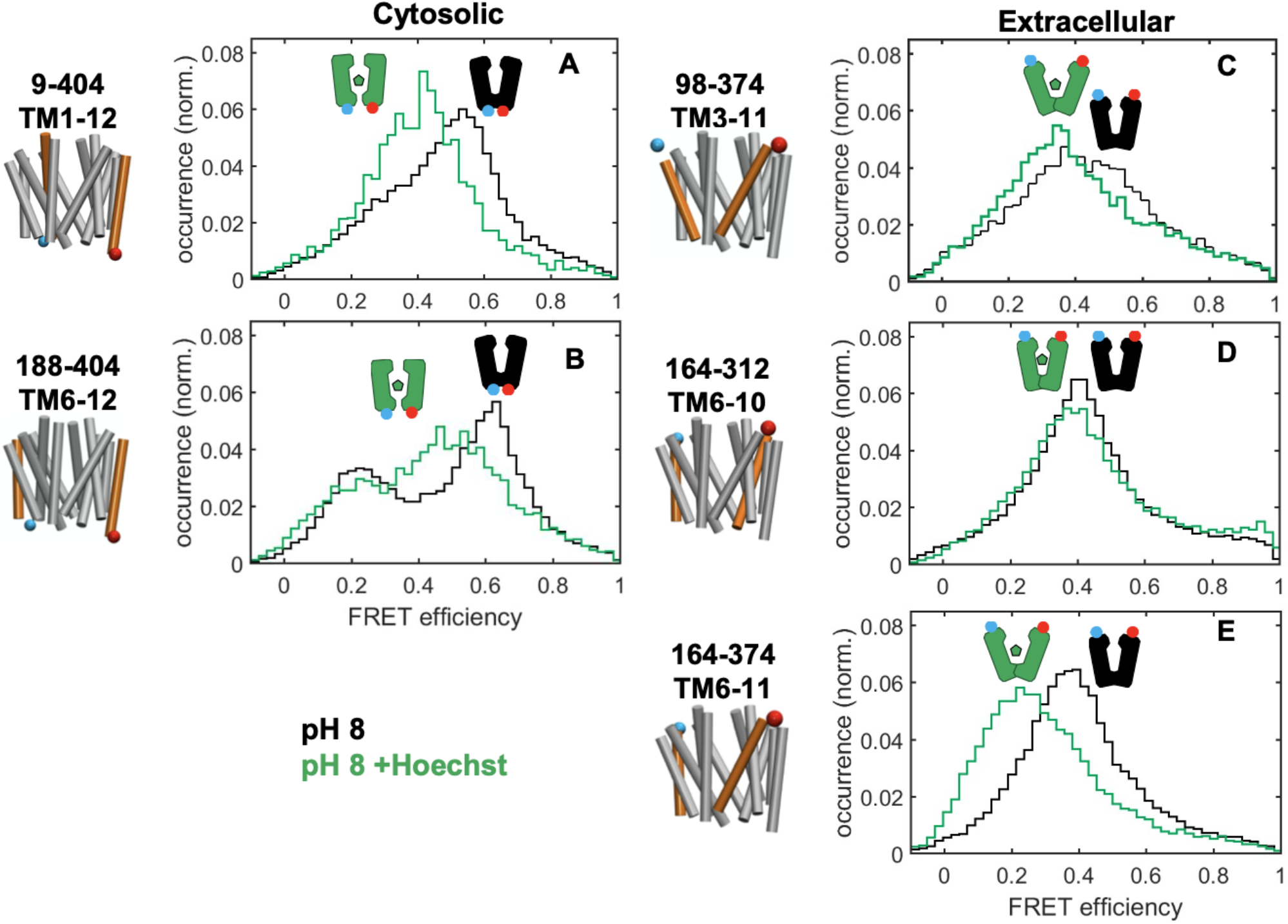
Modulation of the conformational dynamics of LmrP with Hoechst. Left: cartoon representations of the two constructs probing the cytosolic side and the three constructs probing the extracellular side. The helices carrying the probes (in blue (donor) and red (acceptor)) are highlighted in orange. **A-E:** For each mutant measured at pH 8 and at pH 8+Hoechst, the FRET histograms are shown in black and green respectively.

### Opposing effects of Hoechst binding on structural dynamics at the cytosolic and extracellular side

We used photon distribution analysis (PDA) to derive the absolute inter-probe distance distributions from E_FRET_ measurements as well as possible kinetic rate constants for conformer interconversion [25,26]. Specifically, we developed a sequential three-state PDA model to adequately describe the experimental data, as outlined in detail in the materials and methods section.

For the cytosolic distance reporters measured at pH 8 (Fig. 4, Fig. S4 and Table S1), this model included the two states observed in Fig. 2A-B, with mean inter-probe distances ^~^49 Å for the inward-closed and ^~^65 Å for the inward-open state. Interestingly, also a third, very minor ^~^36Å-state was observed that cannot be explained by the outward-open crystal structure. For this minor state, no aberrant donor or acceptor dye fluorescence lifetime, anisotropy or stoichiometry parameters were also observed, ruling out any experimental artifact (Fig. S2). This minor state thus corresponds to a hitherto unidentified ‘inward-very-closed’ state. In terms of interconversion kinetics between states at the cytosolic side, no interconversion of the inward-closed state with the inward-very-closed state was observed for any of the experiments at the cytosolic side (Fig. S5 and Table S1). Between the inward-open and inward-closed state, however, the situation was markedly different. In the absence of Hoechst, the opening rate constant *k*_1→2_ and closing rate constant *k*_2→1_ were on the edge (*k* < 0.01 ms^−1^) of what could be accurately determined with the microscope (Fig. 4D and Fig. S4E, black, and Table S1). The conformational equilibrium was tipped in favor of the inward-closed conformation, as is clear both from the ‘equilibrium constant for closing’ *K*_closing_ > 1, calculated as the ratio between closing and opening rates (Fig. 4D inset and Table S1), and the experimental distance distribution histograms (Fig. 4C and Fig. S4C). In the presence of Hoechst, on the contrary, fast and frequent transitions (with *k*_1→2_ and *k*_2→1_ >0.4 ms^−1^) between the two states were clearly observed (Fig. 4D and Fig. S4E, green, and Table S1), while the steady state distribution between states was almost invariant ((Fig. 4D inset and Table S1), despite of the increased dynamics and thus agrees with the Hoechst-bound crystal structure of the outward-open state. Therefore, at the cytosolic side, the key effect of Hoechst binding is a large increase in conformational interconversion rate rather than an overall conformational population shift.

**Fig.4.**
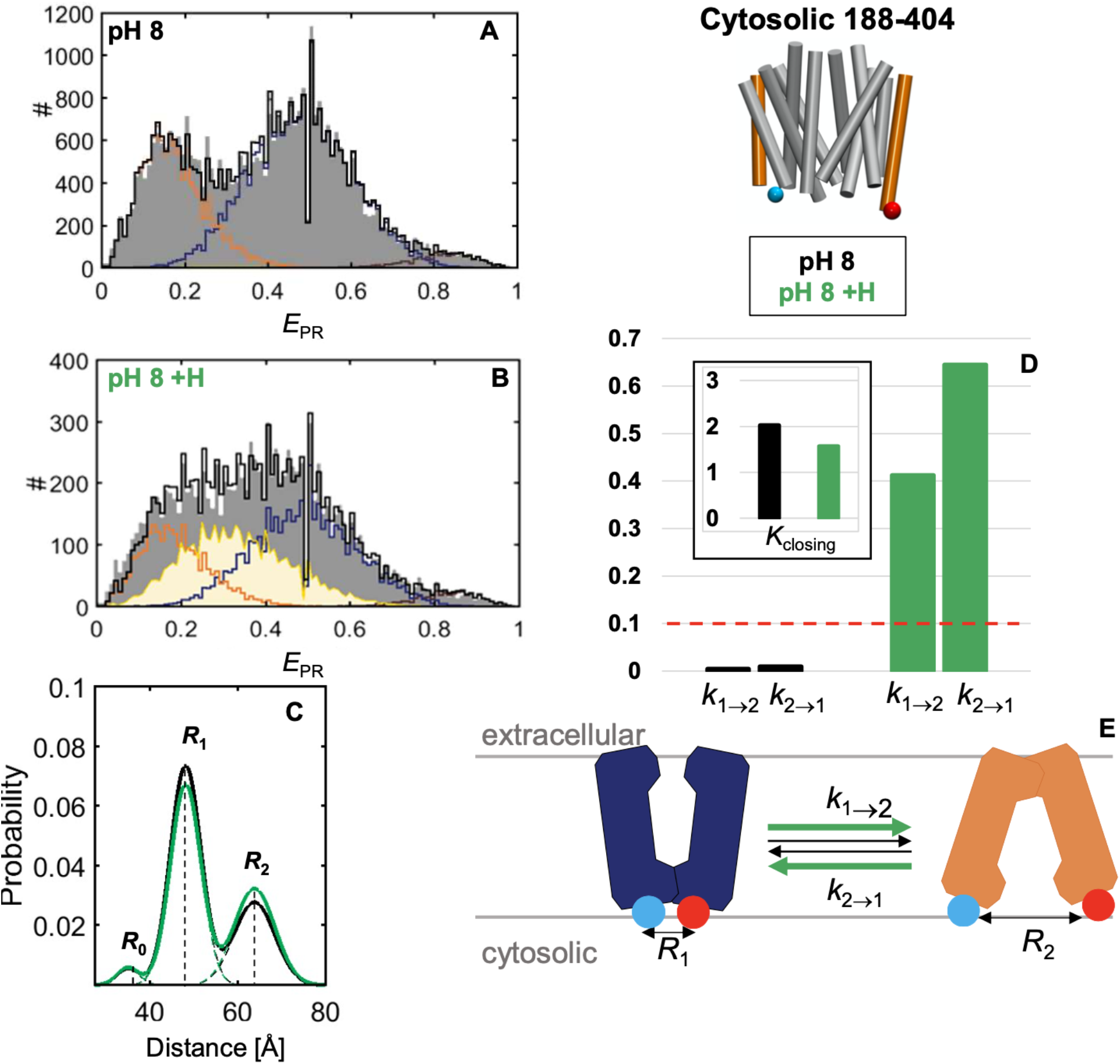
Effect of Hoechst binding on the transition on the cytosolic side. **A-B**: Experimental *E*_PR_ histograms (gray bars) of LmrP 188C-404C burst data rebinned in 1-ms bins and acquired at pH 8 in (**A**) absence or (**B**) presence of Hoechst. The total PDA model (black stairs) is a sum of the inwardopen-state bins (orange stairs), inward-closed-state bins (blue stairs), inward-very-closed-state bins (brown stairs) and bins describing state-interconverting molecules (yellow highlighted area). **C**: The probability density plots of the PDA data from panels A (black solid line) and B (green solid line) are a sum of the underlying components (dashed lines), and are characterized by the distances *R*_0_, *R*_1_ and *R*_2_. **D**: Bar charts of the opening (*k*_1→2_) and closing (*k*_2→1_) rate constants (in ms^−1^) (black = apo, green = Hoechst). The dashed red line is the rate constant below which interconversion kinetics cannot be determined accurately. The inset represents the equilibrium constant for closing. **E.** Cartoon depicting the conformational interconversion kinetics scheme at the cytosolic side (blue = inward-closed, orange = inward-open).

We also used PDA analysis to render distance and kinetic information for the extracellular distance reporters. Three states were observed, an outward-closed (orange *E*_PR_ state in Fig. 5A-B), an outward-open (blue E_PR_ state in Fig. 5A-B) and an outward extra-open state stabilized by Hoechst (purple *E*_PR_ state in Fig. 5A-B). Using this model, fast and frequent transitions were observed in the absence of substrate between all states (Fig. 5D,E and Table S2), with the outward-closed/open equilibrium tipped towards the open state (*K*_closing_ < 1) (Fig. 5C,D/inset, black and Table S2) and the outward-open/very-open equilibrium also tipped towards the open state (*K*_closing_ > 1) (Fig. 5C,E/inset, black and Table S2). In the presence of Hoechst, the frequency of the interconversion decreased (Fig. 5D, green, and Table S2), while again the equilibrium distribution remained similar (Fig. 5C,D/inset,E/inset, green and Table S2). Similar, though not completely identical observations were done for the other extracelullar mutants (Fig. S4F-O). In summary, at the extracellular side the equilibrium is tipped considerably in favor of the outward-open state, and Hoechst decreased the frequency of the transitions between all three observed conformational states.

**Fig.5.**
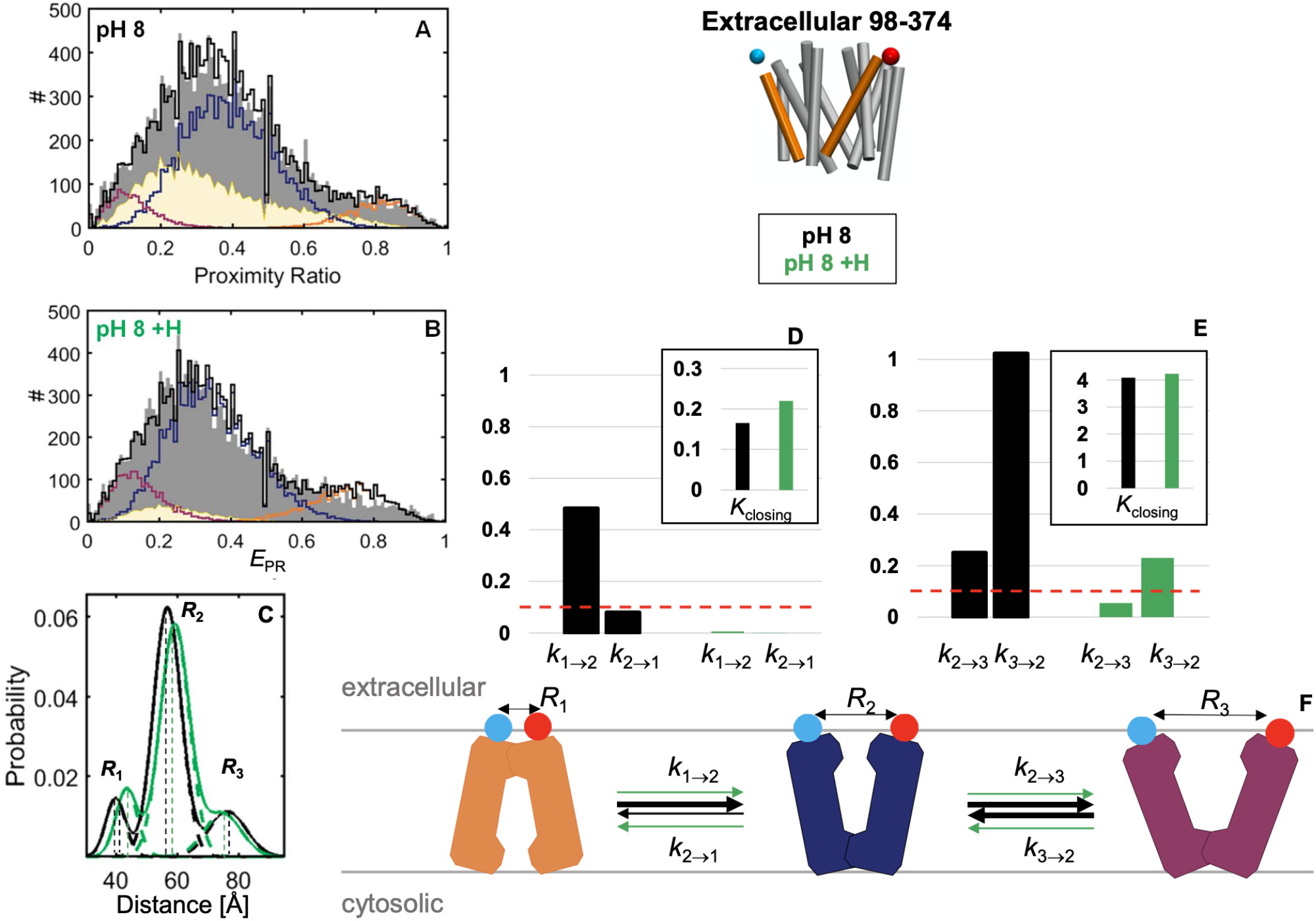
Effect of Hoechst binding on the transition on the extracellular side. **A-B**: Experimental *E*_PR_ histograms (gray bars) of LmrP 98C-374C burst data rebinned in 1-ms bins and acquired at pH 8 in (**A**) absence or (**B**) presence of Hoechst. The total PDA model (black stairs) is a sum of the outward-closed-state bins (orange stairs), outward-open-state bins (blue stairs), extra-open-state bins (purple stairs) and bins describing state-interconverting molecules (yellow highlighted area). **C**: The probability density plots of the PDA data from panels A (black solid line) and B (green solid line) are a sum of the underlying components (dashed lines), and are characterized by the distances *R*_1_, *R*_2_ and *R*_3_. **D**: Bar charts of the opening (*k*_1→2_) and closing (*k*_2→1_) rate constants (in ms^−1^) (black = apo, green = Hoechst). **E**: Bar charts of the extra-opening (*k*_2→3_) and closing (*k*_3→2_) rate constants (in ms^−1^) (black = apo, green = Hoechst). The dashed red lines are the rate constant below which interconversion kinetics cannot be determined accurately. The insets represent the equilibrium constant for closing. **F.** Cartoon depicting the conformational interconversion kinetics scheme at the cytosolic side (orange = outward-closed, blue = outward-open, purple = outward-extra-open).

### Different ligands modulate LmrP conformational dynamics differently

We next assessed whether the observed effect of Hoechst on the LmrP conformational kinetics was universal or substrate-specific by performing similar measurements with another substrate of LmrP, the macrolide roxithromycin. In the presence of roxithromycin, the cytosolic distance reporters adopted two distinct *E* states (Fig. 6D and Fig. S6A, blue). Like with Hoechst, roxithromycin binding led to an increased transition rate while barely affecting *K*_closing_ (Fig. 6E/inset and Fig. S6F/inset, blue). LmrP therefore mostly adopts an inward-closed conformation in all conditions tested. However, the proportion of proteins in the inward-open state increased slightly with roxithromycin (Fig. 6C and S6D), and to a lesser extent, with Hoechst (Fig. 4C and S4C) compared to pH 8 apo.

**Fig.6:**
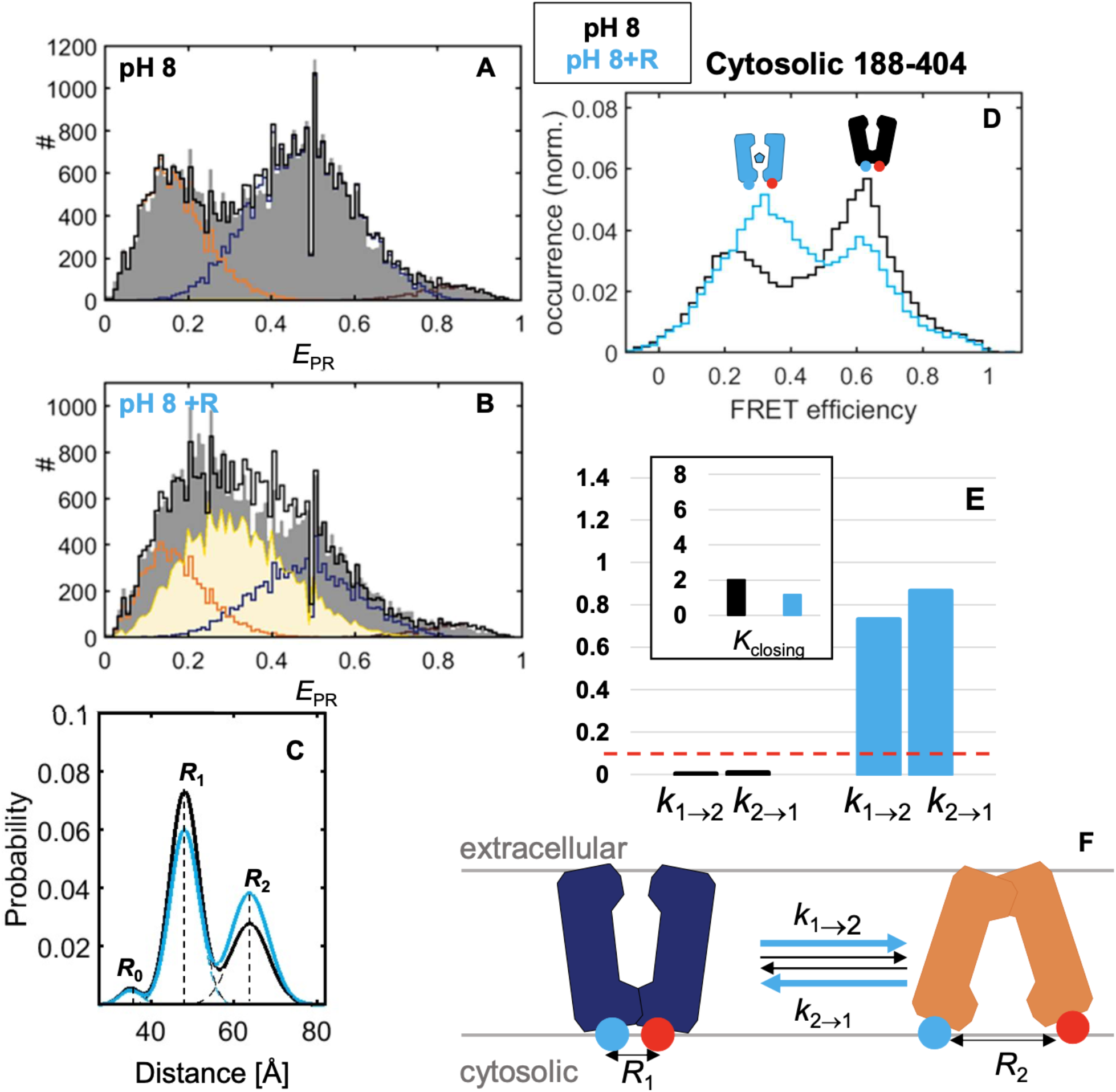
Effect of roxithromycin binding on the transition on the cytosolic side. **A-B**: Experimental *E*_PR_ histograms (gray bars) of LmrP 188C-404C burst data rebinned in 1-ms bins and acquired at pH 8 in (**A**) absence or (**B**) presence of roxithromycin. The total PDA model (black stairs) is a sum of the inward-open-state bins (orange stairs), inward-closed-state bins (blue stairs), inward-very-closed-state bins (brown stairs) and bins describing state-interconverting molecules (yellow highlighted area). **C**: The probability density plots of the PDA data from panels A (black solid line) and B (light blue solid line) are a sum of the underlying components (dashed lines), and are characterized by the distances *R*_0_, *R*_1_ and *R*_2_. **D:** *E* histograms at pH 8 (black) and with roxithromycin (light blue) for LmrP 188C-404C. **E**: Bar charts of the opening (*k*_1→2_) and closing (*k*_2→1_) rate constants (in ms^−1^) (black = apo, light blue = roxithromycin). The dashed red line is the rate constant below which interconversion kinetics cannot be determined accurately. The inset represents the equilibrium constant for closing. **F.** Cartoon depicting the conformational interconversion kinetics scheme at the cytosolic side (blue = inward-closed, orange = inward-open).

On the extracellular side, roxithromycin had little effect on the measured *E* (Fig. 7D and Fig. S6G) and on the proportion of interconverting proteins (Fig. 7A-B and Fig. S6H-I). However, for one of two tested distance reporters, the frequency of the transition between the outward-closed and outward-open states increased with roxithromycin (Fig. 7E), while it decreased between the outward-open and the extra-open state (Fig. 7F). *K*_closing_ was however barely affected in both cases (Fig. 7E-F). As a result, and contrary to Hoechst, roxithromycin did not stabilize an extra-open state of the protein. The outward-open state remained the most populated one with and without roxithromycin, for both extracellular distance reporters. Altogether, while roxithromycin and Hoechst modified the interconversion kinetics similarly on the cytosolic side, they have different effects on the extracellular side of the protein.

**Fig.7:**
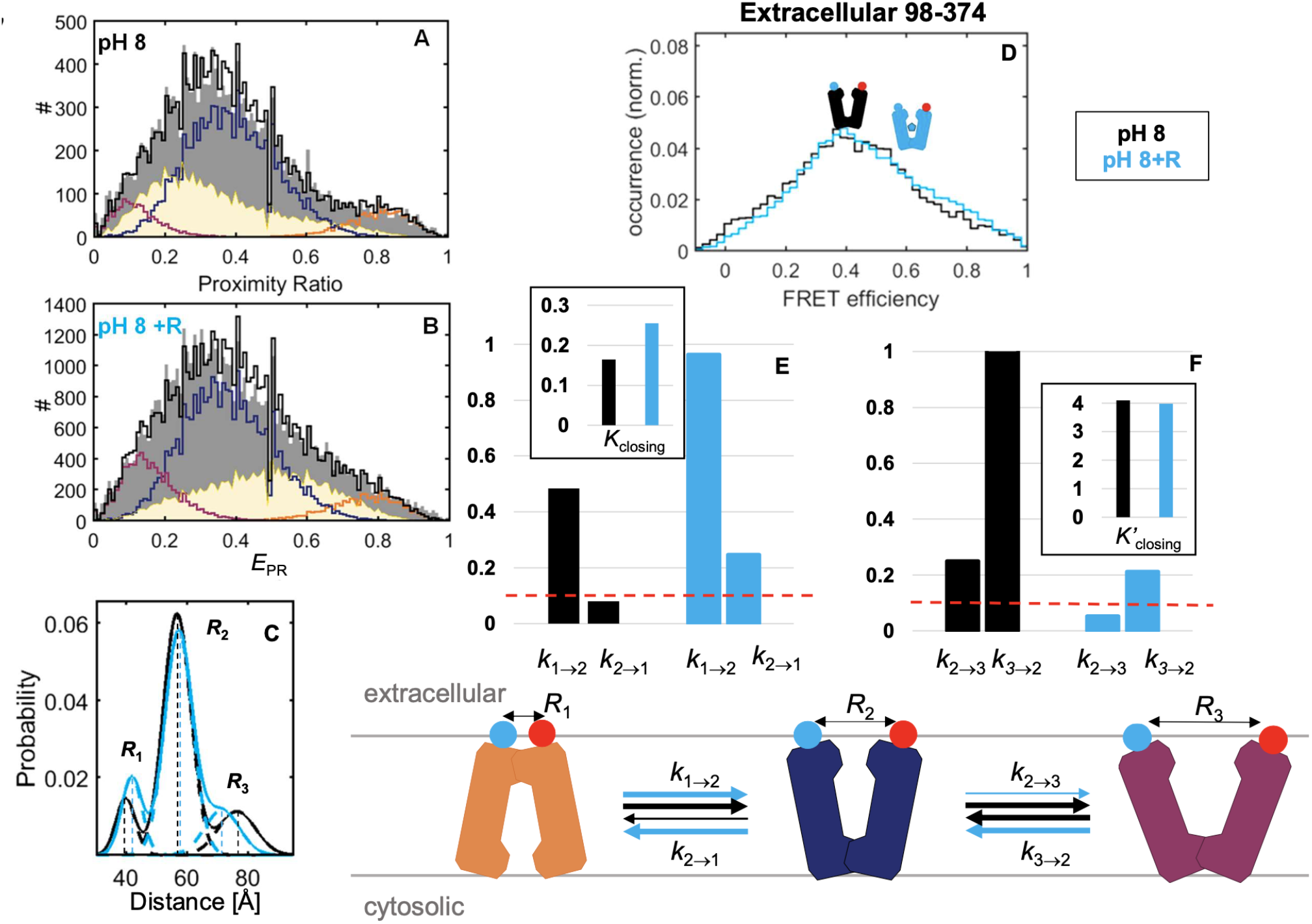
Effect of roxithromycin on the transitions on the extracellular side. **A-B**: Experimental *E*_PR_ histograms (gray bars) of LmrP 98C-374C burst data rebinned in 1-ms bins and acquired at pH 8 in (**A**) absence or (**B**) presence of roxithromycin. The total PDA model (black stairs) is a sum of the outward-closed-state bins (orange stairs), outward-open-state bins (blue stairs), extra-open-state bins (purple stairs) and bins describing state-interconverting molecules (yellow highlighted area). **C**: The probability density plots of the PDA data from panels A (black solid line) and B (light blue solid line) are a sum of the underlying components (dashed lines), and are characterized by the distances *R*_1_, *R*_2_ and *R*3. **D:** *E* histograms at pH 8 (black) and with roxithromycin (light blue) for LmrP 98C-374C. **E**: Bar charts of the opening (*k*_1→2_) and closing (*k*_2→1_) rate constants (in ms^−1^) (black = apo, light blue = roxithromycin). **F**: Bar charts of the extra-opening (*k*_2→3_) and closing (*k*_3→2_) rate constants (in ms^−1^) for all extracellular reporter mutants (black = apo, light blue = roxithromycin). The dashed red line is the rate constant below which interconversion kinetics cannot be determined accurately. The insets represent the equilibrium constant for closing. **G.** Cartoon depicting the conformational interconversion kinetics scheme at the cytosolic side (orange = outward-closed, blue = outward-open, purple = outward-extra-open).

## III. Discussion

Our study shows that smFRET can characterize the conformational equilibrium and interconversion kinetics of the membrane transporter LmrP. We observed that the protein mostly adopts two conformations, inwardopen and outward-open, in an equilibrium modulated by pH, substrate binding and a specific mutation. While such smFRET measurements were performed in solution at room temperature and pM concentration of protein, our data agrees with previous DEER measurements, even though these are performed at μM concentration in the frozen state[12].

While the equilibrium between outward-open and inward-open states on MFS transporters is typically described via the rocker-switch model [27], our data shows that the cytosolic and extracellular sides of the protein are in fact uncoupled. This is particularly clear in the presence of Hoechst where the opening and closing rate constants of the protein are increased on the cytosolic side and decreased on the extracellular side. Furthermore, our data not only identifies conformational heterogeneity between the intra- and the extracellular sides of the proteins but also between different helices on a given side. For instance, the distance reporter 164C-312C seems to be less sensitive to distance and kinetics changes than the others. This indicates that the protein is highly flexible and does not perform its transport mechanism simply through rigid body motions.

Using PDA, we have shown that ligands affect only slightly the steady state distribution protein conformations, but rather modulate the interconformeric transition rates. Interestingly, we also observed that different ligands affect the transition kinetics differently. While roxithromycin had the same effect than Hoechst on the interconversion kinetics at the cytosolic side, at the extracellular side, and contrary to Hoechst, roxithromycin increased the dynamics of the transitions between the outward-closed-open/states. The impact of increased dynamics at the cytosolic side is intriguing and could suggest that Hoechst and roxithromycin may lower the energy barrier between the inward-closed and inward-open conformations. We recently proposed that the ability of LmrP to bind structurally diverse ligands does not rely on structural adaptation but is rather enabled by the presence of an embedded lipid inside the binding pocket, that provides a conformational malleable hydrophobic environment. This study, however, shows that the kinetic behavior of the protein can vary with different ligands. This could suggest differences in the actual transport mechanism across the spectrum of possible substrates.

The binding of an electroneutral substrate to the MDR transporter MdfA has been recently shown to stabilize an outward-closed conformation and to decrease the frequency of the transition on the extracellular side, while the binding of a cationic substrate stabilized an outward-open conformation without affecting the interconversion kinetics [28]. Different ligands also seem to modulate LmrP kinetics differently. Other ligands should therefore be tested to further characterize their particular effects.

While our study indicates that the smFRET/MFD-PIE approach is a good method to characterize conformational transition of membrane transporters, one limitation is that is tracks relatively fast conformational changes (<10ms) due to the limited dwell time of the molecules in the confocal volumes. As a comparison, the frequency of the transition of the surface-immobilized MDR transporter MdfA has been recently characterized with total internal reflection fluorescence (TIRF)-smFRET at 0.2-0.3 s^−1^ for the extracellular side [28]. A few years before, the opening/closing rates of the MFS importer LacY have also been characterized by measuring the (un)quenching by a Trp residue of the fluorescence of a bimane probe attached to a Cys residue. The spontaneous opening of the periplasmic side happened at 20-30 s^−1^, while the opening/closing rate of the cytosolic side was slower (around 4-8 s^−1^) [29]. Clearly, transporters exhibit conformational changes on different timescale. An extensive characterization of the proteins with different methods giving access to different timescales is therefore recommended. TIRF-smFRET on surface-immobilized proteins could be used to characterize LmrP dynamics on the ms-s timescale [30], while correlation analysis, in particular filtered fluorescence correlation spectroscopy (fFCS), could be used to study the dynamics on the microsecond timescale [31]. Moreover, future work should also address differences between different FRET pairs, each one probing a different portion of the protein. Finally, we should stress that measurements were performed in detergent micelles and it will be interesting to investigate whether the presence of a lipid bilayer (i.e. in a proteoliposome or a nanodisc), will affect the conformational dynamics (as in [32] for instance).

## V. Methods

Hoechst 33342 is referred to as Hoechst in the article.

### Bacterial strains, plasmids and growth conditions

LmrP was expressed in the *L. lactis* NZ9000 strain that is deleted from the known ABC multidrug transporters LmrA and LmrCD. We used the in-home pHLP5 plasmid, a derivative of the *L. lactis* expression vector pNZ8048 carrying the *lmrP* gene coding for the C-terminally 6 Histidine-tagged LmrP [35]. Briefly, cells were grown at 30 °C in M17 broth supplemented with 0.5% (w/v) glucose and 5 μg/mL chloramphenicol until the OD_660_ reaches 0.8-0.9. The overexpression of LmrP was then induced by addition of 1:250 dilution of the supernatant of the nisin producing L. lactis strain NZ9700. After 2h of induction at 30 °C, cells were harvested by centrifugation at 7000 × g.

### Design and construction of the mutants

For each distance reporter, two residues were selected to be mutated into cysteines based on the structure of LmrP obtained by X-ray crystallography in the outward-open conformation (PDB 601Z) as well as on the model of the inward-open conformation obtained by inverted-topology repeats by our collaborator José D. Faraldo-Gómez. Cysteines were placed at the extra-or intracellular extremities of chosen TMs while avoiding mutating conserved residues. The expected FRET efficiencies were estimated by simulating the fluorescent probes’ accessible volume [33] so that they are separated by a distance within the Förster radius, 56.8 Å in this case[36]. The accessible volume (AV) clouds were simulated using the parameters given for ATTO488 and Alexa647 in [36].

The plasmid template was first methylated using the dam methyltransferase (NEB) following the manufacturer recommendations and as described in [37]. Mutations were introduced one by one by site-directed mutagenesis using the *Pfu* DNA polymerase (Promega) in the in-home pHLP5 plasmid in which the only endogenous Cys270 had been previously replaced by an alanine. The primers were designed as described in [38]. After transformation, plasmid DNA was extracted and verified by sequencing.

### Preparation of inside-out membrane vesicles

As described in [35], cells were washed in 50 mM HEPES pH 7 and resuspended (10 mL for each L of culture) in the same buffer supplemented with 7 mg/mL of lysozyme from chicken egg whites, 10 μg/mL Deoxyribonuclease I from bovine pancreas, 10 mM MgSO4 and 50 mM Dithiothreitol (DTT). After a 1h30 incubation at 30 °C, cells were homogenized with a tissue grinder Potter-Elvehjem type and broken by four passes at 1000–1500 bars using a high-pressure homogenizer. Cell debris and undisrupted cells were subsequently eliminated by centrifugation at 17 000 × g. Inside-out membrane vesicles were then isolated by ultracentrifugation of the supernatant at 100 000 × g for 2h at 4 °C. Membranes were resuspended in a pH7 buffer composed of 100 mM HEPES, 300 mM NaCl, 20% (w/v) glycerol (5 mL buffer/L culture) and 1 mM DTT.

### Transport assays

The transport activity of LmrP mutants was assessed by measuring Hoechst fluorescence (EX 355 nm, EM 457 nm) decay over time at 30 °C, based on previously published protocols [12,35,39]. Briefly, inside-out membrane vesicles of LmrP-expressing cells (^~^ 1 mg proteins) were diluted in transport buffer (50 mM HEPES, 2 mM MgCl_2_, 300 mM KCl, pH 7.4) with 0.3μM of Hoechst (Invitrogen). Addition of 3 mM of ATP-Na2 in 50 mM HEPES pH 6.8 buffer supplemented with 250 mM MgSO_4_ allows the generation of a proton gradient by activating the endogenous F_0_F_1_-ATPase. Active extrusion of Hoechst by LmrP is detected as a decrease of fluorescence is measured. Addition of 1μM of the ionophore nigericin dismisses the proton gradient and Hoechst is no longer actively extruded by LmrP causing the fluorescence to increase. Raw data were normalized for fluorescence intensities relative to the initial fluorescence prior to ATP addition.

### LmrP purification & labelling for smFRET

Inside-out membrane vesicles were solubilized in a solution of 2.5% (w/V) n-dodecyl- β – D-maltoside (β-DDM) in water with 1 mM DTT. After 1.5 h incubation on a rotating wheel at 4 °C, the insoluble part was removed by ultracentrifugation at 100 000 × g for 1 h. The supernatant was batch-incubated with Ni^2+^-nitrilotriacetate affinity resin (Ni-NTA; Qiagen; 500 μL resin/L of culture) with 10 mM imidazole at 4 °C for 2 h. The resin was pre-equilibrated with buffer A (50 mM HEPES, 150 mM NaCl, 10% (w/V) glycerol, 20 mM imidazole and 0.05% (w/V) β-DDM, pH 7). After the incubation, the slurry was transferred into a polypropylene column, the flow-through discarded, and the resin was washed with 8 CV of buffer A, supplemented with 0,5 mM DTT. LmrP was then eluted by stepwise additions of buffer B (buffer A with 250 mM imidazole and 0,5 mM DTT). The more concentrated fractions were pooled together. Imidazole and DTT were removed just prior to labelling using a desalting column (Disposable PD10 desalting columns, GE Healthcare) previously equilibrated with labelling buffer (50 mM HEPES, 150 mM NaCl and 0.02% (w/V) β-DDM, pH 7.5) that was degassed in vacuum/N_2_ under continuous stirring. LmrP was concentrated to approximately 100 μM using a 50 kDa MWCO concentrator (Amicon, GE Healthcare) and then labelled with the donor ATTO488 (ATTO 488 maleimide, ATTO-TEC GmbH) and the acceptor Alexa647 (Alexa Fluor 647 maleimide, Life Technologies Europe BV). 250 μM Alexa647 and 100 μM ATTO488 were mixed in the degassed Labelling buffer and 50 μM of LmrP was added for 2 h at RT. The majority of unreacted dye was removed on a desalting column and the labelled protein was run on an SDX-200 10/300 GL (GE Healthcare) size exclusion chromatography column in SEC buffer (50 mM HEPES pH 7 150 mM NaCl, 10% (w/v) glycerol and 0.02% (w/v) β-DDM). The efficiency of the labelling was checked via UV-visible absorbance at 280 nm, 501 nm and 650 nm to determine the amount of LmrP, ATTO488 and Alexa647 respectively, after correcting the 280-nm peak for the dyes’ absorbance.

### smFRET sample preparation

Measurement buffers were filtered through a 0.22 μM filter and supplemented with 0.01% (w/V) β-DDM (Anatrace) just before use. pH 8 measurement buffer: 50 mM HEPES (CellPURE, Fisher Bioreagents), 150 mM NaCl (TraceSELECT, Honeywell Fluka). pH 5 measurement buffer: 25 mM MES (Hydrate, Sigma), 25 mM acetate (for analysis, Merck), 150 mM NaCl (TraceSELECT, Honeywell Fluka).

Experiments were performed on freshly labelled LmrP (not frozen). The protein was diluted right before each measurement in the desired measurement buffer, such that the probability of having >1 molecule in the focus was negligible (which corresponded to an average count rate per detector of 3–4 kHz, or ^~^50pM protein concentration). 30 μL of the sample was put on the coverslip of an 8-well chamber cuvette (Nunc Lab-Tek Chambered Coverglass, Thermo Fisher Scientific), previously coated with 30 μL bovine serum albumin (BSA) (1 mg/mL in PBS) and washed twice with the sample before starting the measurement. The sample holder was closed with a lid to avoid evaporation during the measurements as well as contamination.

If LmrP was measured with a substrate, LmrP was first incubated with this substrate for 30 minutes on ice. (^~^0.5-1 μM LmrP and 1 mM roxithromycin). For the finale dilution before the measurement, roxithromycin was added at 100μM. For Hoechst, since the substrate is also fluorescent, LmrP was incubated with 100μM of Hoechst and measured with 3 μM of Hoechst to avoid any additional background in the experiments.

Background profiles (needed for calculating E and S parameters, and for lifetime and PDA analysis) were prepared similarly, but without protein and were recorded twice for five minutes.

### smFRET data recording

per sample, because the protein was unstable upon longer measurements, samples were recorded at least three times 30 minutes at room temperature (22 °C). Fresh dilutions were performed for each measurement. Measurements were performed on a homebuilt multiparameter fluorescence detection microscope with pulsed interleaved excitation (MFD-PIE) with pulsed interleaved excitation (MFD-PIE) as established^50^, with minor modifications. Emission from a pulsed 483-nm laser diode (LDH-P-C-470, PicoQuant) was cleaned up (Chroma ET485/20x, F49-482; AHF analysentechnik AG), emission from a 635-nm laser diode (LDH-P-C-635B, PicoQuant) was cleaned up (Chroma z635/10x, PicoQuant), and both lasers were alternated at 26.67 MHz (PDL 828 Sepia II, PicoQuant), delayed ^~^18 ns with respect to each other, and combined with a 483-nm re-flecting dichroic mirror in a single-mode optical fiber (coupler, 60FC-4-RGBV11-47; fiber, PMC-400Si-2.6-NA012-3-APC-150-P, Schäfter + Kirchhoff GmbH). After collimation (60FC-L-4-RGBV11-47, SuK GmbH), the linear polarization was cleaned up (CODIXX VIS-600-BC-W01, F22-601; AHF analysentechnik AG), and the light (100 mW of 483-nm light and 50 mW of 635-nm light) was reflected into the back port of the microscope (IX70, Olympus Belgium NV) and upward [3-mm-thick full-reflective Ag mirror, F21-005 (AHF) mounted in a total internal reflection fluorescence filter cube for BX2/IX2, F91-960; AHF analysentechnik AG] to the objective (UPLSAPO-60XW, Olympus). Sample emission was transmitted through a 3-mm-thick excitation polychroic mirror (Chroma zt470-488/640rpc, F58-PQ08; AHF anal-ysentechnik AG), focused through a 75-mm pinhole (P75S, Thorlabs) with an achromatic lens (AC254-200-A-ML, Thorlabs), collimated again (AC254-50-A-ML, Thorlabs), and spectrally split (Chroma T560lpxr, F48-559; AHF analysentechnik AG). The blue range was filtered (Chroma ET525/50m, F47-525, AHF analysentechnik AG), and polarization was split (PBS251, Thorlabs). The red range was also filtered (Chroma ET705/100m, AHF analysentechnik AG), and polarization was split (PBS252, Thorlabs). Photons were detected on four avalanche photodiodes (PerkinElmer or EG&G SPCM-AQR12/14, or Laser Components COUNT BLUE), which were connected to a time-correlated single-photon counting (TCSPC) device (SPC-630, Becker & Hickl GmbH) over a router (HRT-82, Becker & Hickl) and power supply (DSN 102, PicoQuant). Signals were stored in 12-bit first-in-first-out (FIFO) files. Microscope alignment was carried out using fluorescence correlation spectroscopy (FCS) on freely diffusing ATTO 488-CA and ATTO 655-CA (ATTO-TEC) and by connecting the detectors to a hardware correlator (ALV-5000/EPP) over a power splitter (PSM50/51, PicoQuant) for alignment by real-time FCS. Instrument response functions (IRFs) were recorded one detector at-a-time in a solution of ATTO 488-CA or ATTO 655-CA in near-saturated centrifuged potassium iodide at a 25-kHz average count rate for a total of 25e6 photons. Macrotime-dependent microtime shifting was corrected for two (blue/parallel and red/perpendicular) of four avalanche photodiodes (APDs) using the IRF data as input^51^.

### Single-Molecule Burst Analysis

Data was analyzed with the PAM software^52^ via standard procedures for MFD-PIE smFRET burst analysis^53,54^. Signals from each TCSPC routing channel (corresponding to the individual detectors) were divided in time gates to discern 483-nm excited FRET photons from 635-nm excited acceptor photons. A two-color MFD all-photon burst search algorithm using a 500-ms sliding time window (minimum of 50 photons per burst, minimum of 5 photons per time window) was used to identify single donor- and/or acceptor–labeled molecules in the fluorescence traces. Double labelled single molecules were selected from the raw burst data using a kernel density estimator (ALEX-2CDE < 15), that also excluded other artefacts. Bursts with a donor lifetime <0.011 or >5 ns and with and acceptor lifetime <0.011 or >3 ns were omitted from the data. Sparse slow-diffusing aggregates were removed from the data by excluding bursts exhibiting a burst duration >30 ms. Data was corrected in this order to obtain the absolute stoichiometry parameter *S* and absolute FRET efficiency *E*: background subtraction, donor emission crosstalk correction, acceptor direct excitation correction, relative detection efficiency correction. By histogramming *E* versus measurement time, we corroborated that the distribution of *E* was invariant over the duration of the measurement.

MFD-PIE allows for each molecule the determination of the *E*, the labelling stoichiometry, the fluorescence lifetime, and the anisotropy of the donor and the acceptor that are represented in the multi-parameter graphs in Fig S2. The single-burst analysis was performed as described in [36]. Briefly, the absolute burst-averaged FRET efficiency *E* was calculated with:

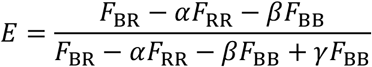

Where *F*_BR_, *F*_RR_ and *F*_BB_ are the background corrected number of photons detected in the red detection channels after blue excitation, in the red detection channels after red excitation and in the blue detection channels after blue excitation, respectively; *α* is a correction factor for the direct excitation of the acceptor with the 483 nm laser; *β* is a correction factor for the emission crosstalk of the donor in the acceptor channel and *γ* is a correction factor for the relative detection efficiency of the donor and acceptor [17].

The corrected stoichiometry (*S*) is given by:

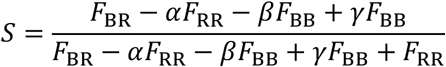

The single-molecule burst-averaged fluorescence lifetimes of FRET donor and acceptor lifetimes were determined using a maximum likelihood estimator approach (MLE) [40]. The static FRET lines were calculated assuming an R_0_ = 56.8 Å, a donor-only lifetime estimated from the experimental data (*S* value> 0.8), and the standard deviation of the Gaussian linker dye distribution of 5.6 Å. The FRET donor (*r*_D_) and acceptor anisotropy (*r*_A_) were calculated from the burst-wise intensities in the different polarization channels. The anisotropy (*r*) versus fluorescence lifetime plots were fitted with a Perrin equation:

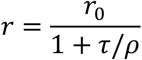

Where *r*_0_ is the fundamental anisotropy (*r*_0_ = 0.4), *τ* is the fluorescence lifetime and *ρ* is the rotational correlation time defined as the average time the dye takes to rotate over 1 radian [41].

### Three-state photon distribution analysis

Photon distribution analysis (PDA) [25,42] provides a statistical analysis of the distribution of photon counts into the donor and acceptor spectral channels. Through this approach, it is possible to separate the contributions of photon shot noise from the physical heterogeneity of the studied system, which may originate either from a distribution of static conformations or from dynamic interconversion between distinct states during the observation time.

Practically, for each FRET data set, raw bursts were re-binned into 1-ms. Binned data was plotted in a raw (uncorrected) FRET efficiency (*E*_PR_) versus uncorrected stoichiometry (*S*_PR_) plot and only keeping the bins with 0.3<*S*_PR_<0.5 efficiently removed burst sections containing complex acceptor photophysics or photobleaching. Furthermore, only bins with at least 10-20 and maximally 250 photons (to reduce calculation time) were used for PDA analysis.

Briefly, for a given FRET efficiency *E*, PDA provides a semi-analytical description of the expected shot-noise limited distribution of the proximity ratio, 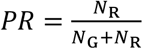, where *N*_G_ and *N*_R_ are the photon counts in the donor and FRET channels, respectively. Under the assumption of a binomial distribution for the FRET process with a success probability equal to the FRET efficiency *E*, the proximity ratio histogram *h^PR^* is described by [25]:

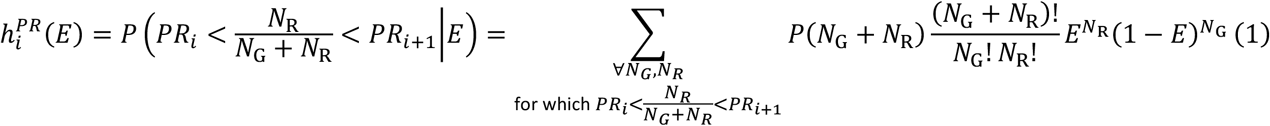

Here, *h_i_*(*E*) is the value of the *i*-th bin of the shot-noise limited histogram defined by the FRET efficiency *E*, *PR_i_* and *PR*_*i*+1_ are the respective bin edges, and *P*(*N*_G_ + *N*_R_) is the empirical distribution of photon counts that determines the shot-noise. *P*(*N*_G_ + *N*_R_) describes the distribution of the number of photons per singlemolecule event (i.e. the burst size distribution) and is obtained from the experiment. Additional corrections for experimental artefacts (spectral crosstalk, direct excitation and background) are accounted for as described in reference [25].

In the case of a distribution of FRET efficiencies *E*, the total shot-noise limited histogram *H* is given by the weighted sum over all contributions:

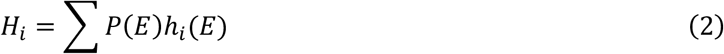

where *P*(*E*) is the probability to observe a single-molecule event with the average FRET efficiency *E*. To account for static or dynamic heterogeneity, one needs to find an expression for the distribution of FRET efficiencies. For static heterogeneity, it is customary to use a Gaussian distribution of distances that is converted into the corresponding FRET efficiency distribution *P*(*E*) [25], although arbitrary distributions or model-free approaches may be applied as well [43].

To describe the effect of dynamics between distinct states with different FRET efficiencies on the proximity ratio histogram, one has to consider the fraction of the time the molecule spends in the different states, given by 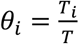 where *T_i_* is the time spent in state *i* and *T* is the observation time. The distribution of the fractional occupancy of the states, *P*(*θ_i_*|*T*, ***K***), depends on the transition rate matrix ***K*** and is known in analytical form only for two-state systems [44,45]. For the description of the dynamics of LmrP, we use a linear three-state dynamic model of the form:

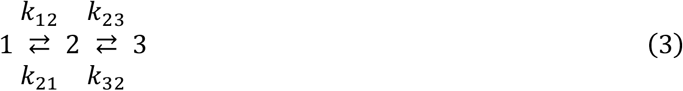

The occupation time distribution *P*(*θ_i_*|*T*, ***K***) is approximated by Monte Carlo simulations using the Gillespie algorithm [46] by simulating the system for many (>100,000) observation time windows, similar to a previously described approach [47]. The average FRET efficiency 〈*E*〉 of a single-molecule event then depends on the fractional occupations of the different states:

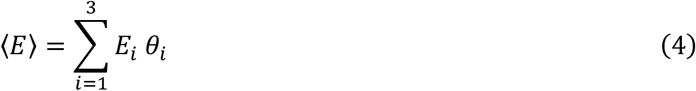

Generally, the different FRET efficiency states may have different brightness, in which case the average FRET efficiency is given by the weighted sum [26]:

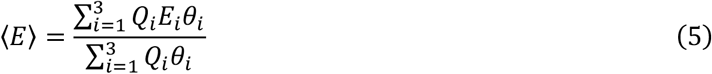

where *Q_i_*, *E_i_* and *T_i_* are the relative brightness, the FRET efficiency and the time spent in state *i*. Through this relation, the occupation time distribution defines the distribution of the FRET efficiency, *P*(*E*) = *P*(〈*E*〉), which is used to construct the shot-noise limited proximity ratio histogram by equations 1 and 2 above.

The seven fit parameters (three FRET efficiencies and four rate constants) were refined using the Nelder-Mead optimization algorithm [48] as implemented in the MATLAB function *fminsearch*. To increase the robustness of the fitting, we exploit the fact that proximity ratio histograms with different observation times *T* can be generated from a single dataset. These histograms show different kinetic signatures due to the different integration times [26] but are described by identical kinetic parameters. In addition, we assumed that the conformational states of LmrP in the presence and absence of ligands were identical and therefore fitted the datasets of LmrP with and without ligands globally with respect to the FRET efficiencies of the three conformational states. For the distance reporter 98C-374C however, a reasonable fit could not be achieved this way, and the distances were separately optimised for each condition (pH 8 apo and with ligands). Simultaneously, the kinetic rates are optimized separately for each condition (± ligand) by globally fitting the histograms generated for four different observation times (T = 0.2, 0.5, 0.75, 1 ms).

We defined the equilibrium constants for closing as:

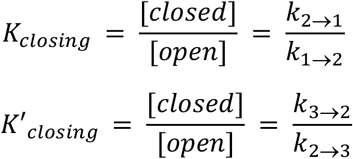

### Software

smFRET data were analyzed using the software package PAM developed by the group of Don C. Lamb (LMU Munich) [49]. A detailed manual can be found at http://pam.readthedocs.io.

AV calculations were done with software developed by the group of Claus Seidel (HHU Düsseldorf) [33], available at http://www.mpc.hhu.de/software/fps.html.

## VII. Acknowledgements

Johan Hofkens is acknowledged for the use of his imaging facilities.

José D. Faraldo-Gómez for the inward-open model of LmrP.

Eva Marting and Niels Vandenberk are thanked for their technical assistance.

## Notes

### Competing Interest Statement

The authors have declared no competing interest.

