## Supporting Information for "Substrate binding modulates the conformational kinetics of the secondary multidrug transporter LmrP"

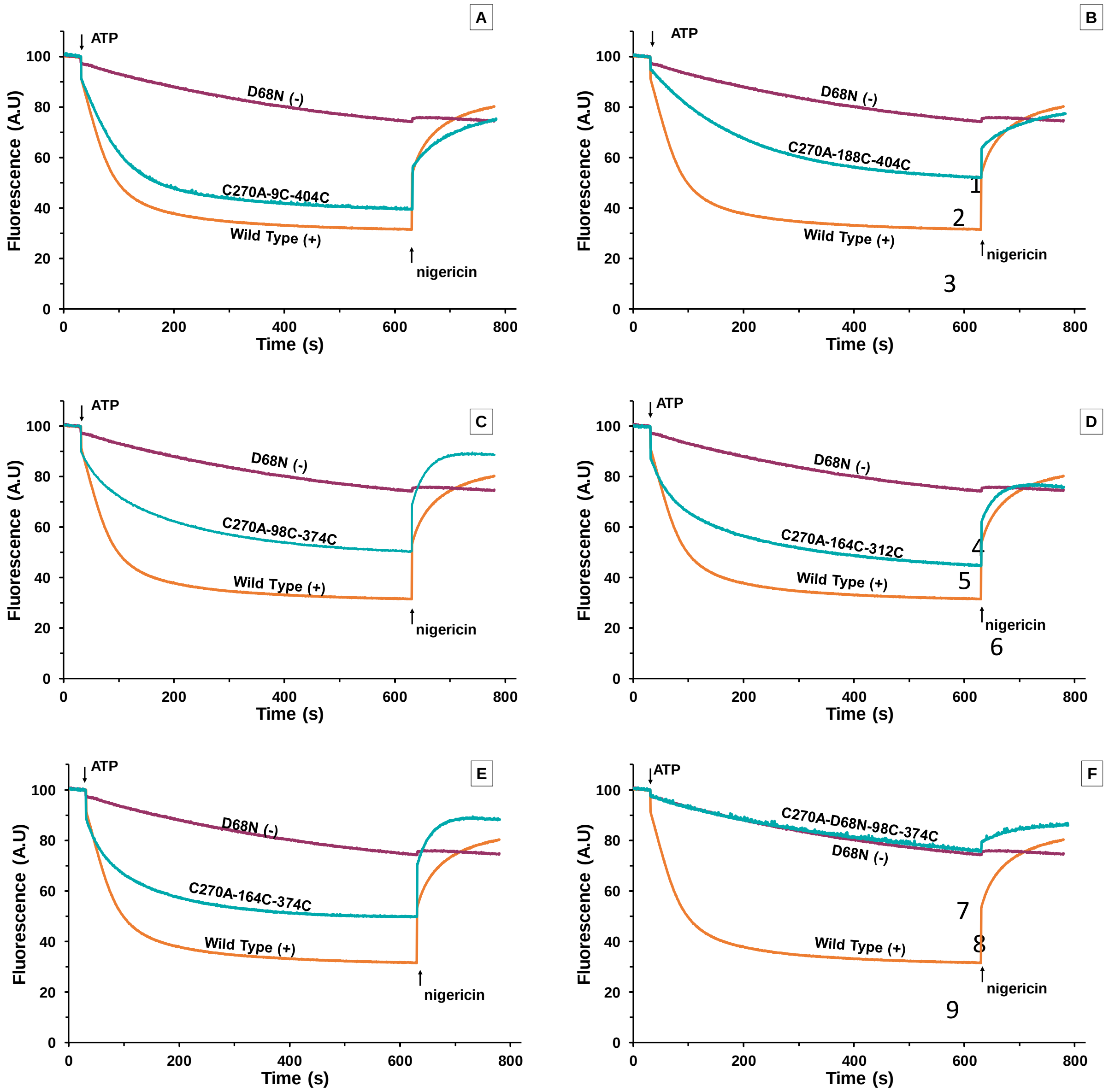


**Fig S.1: Transport activity assay of the double cysteine mutants:**
The extrusion of Hoechst by LmrP in inside-out membrane vesicles is followed by a fluorescence-based assay (Exc: 355nm, Em: 457nm). Values were normalised for fluorescence intensity. ATP energises the endogenous $F_0F_1$-ATPase, generating a transmembrane proton gradient which in turn activates LmrP. The addition of nigericin disrupts the proton gradient and inactivates LmrP. Hoechst is transported by LmrP WT (in orange, positive control) but not by the D68N mutant (in purple, negative control). The double cysteine mutants C270A-9C-404C (**A**, blue), C270A-188C-404C (**B**, blue), C270A-98C-374C (**C**, blue), C270A-164C-312C (**D**, blue) and C270A-164C-374A (**E**, blue) transport Hoechst as well. The mutant C270A-D68N-98C-374C (**F**, blue) does not transport Hoechst.

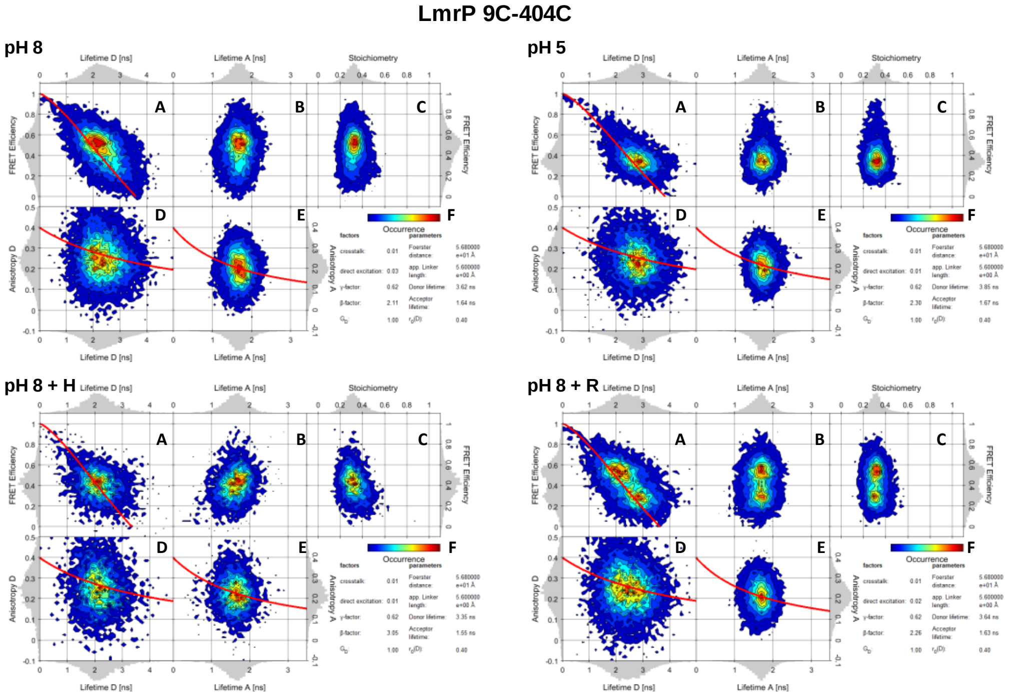
LmrP 9C-404C
pH 8
pH 5
pH 8 + H
pH 8 + R
A
B
C
D
E
F
Lifetime D [ns]
Lifetime A [ns]
Stoichiometry
FRET Efficiency
Anisotropy D
Anisotropy A
Occurrence

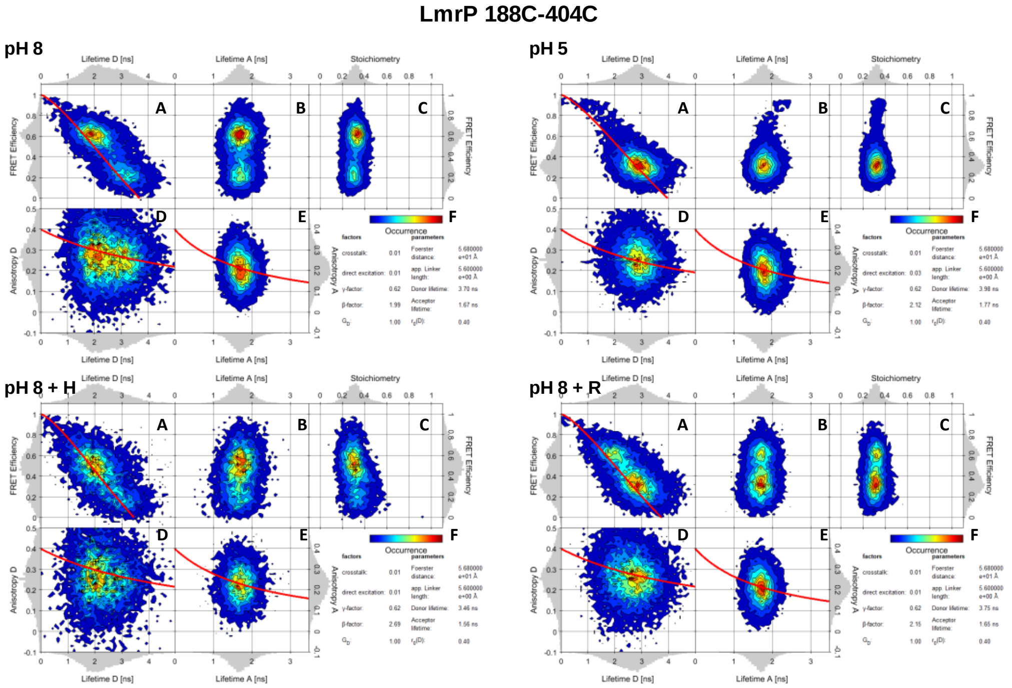
LmrP 188C-404C
pH 8
pH 5
pH 8 + H
pH 8 + R
A
B
C
D
E
F
Lifetime D [ns]
Lifetime A [ns]
Stoichiometry
FRET Efficiency
Anisotropy D
Anisotropy A
Occurrence

**LmrP 98C-374C**

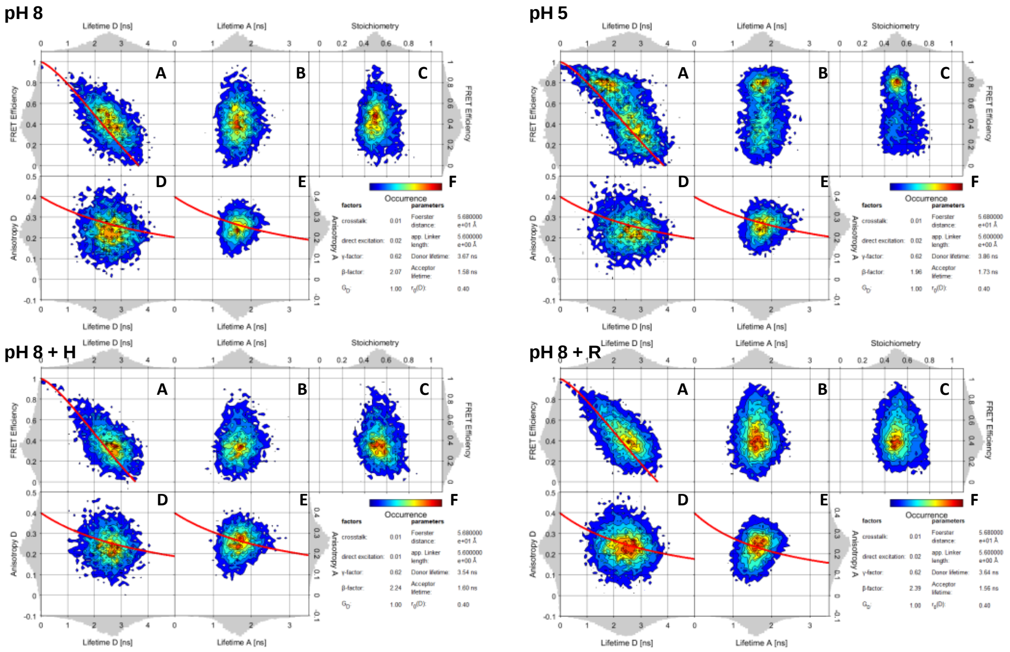


21

**LmrP 164C-312C**

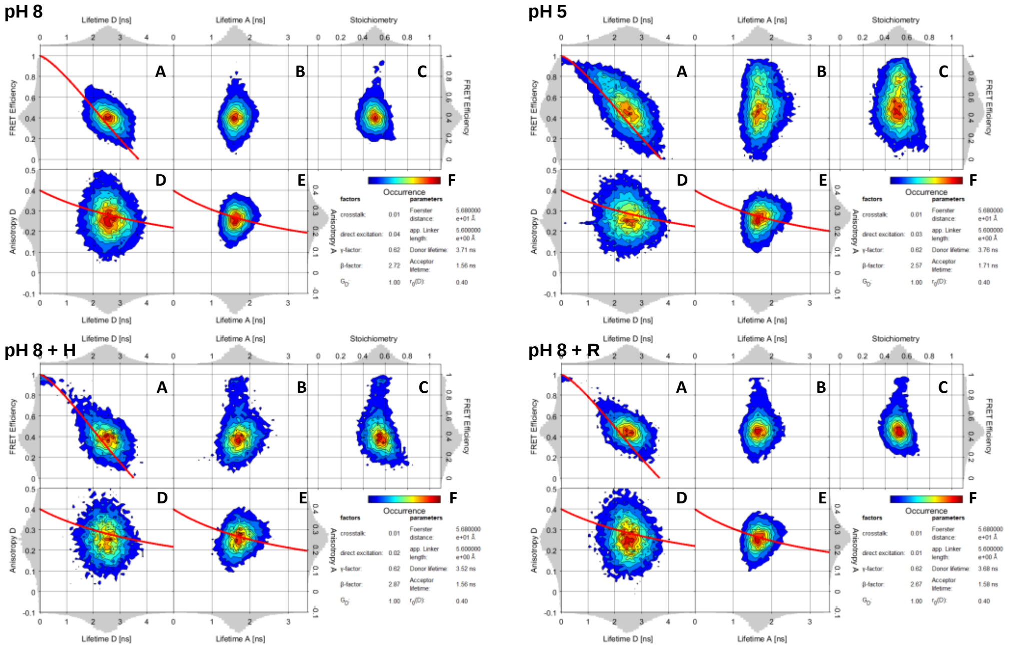


22

**LmrP 164C-374C**

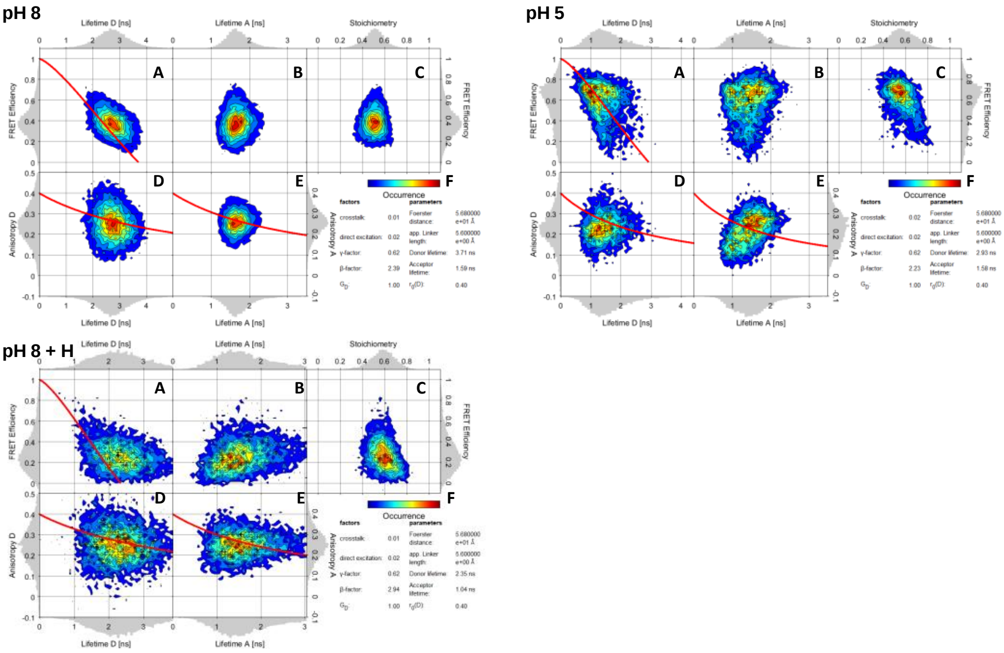


**Fig S.2: Multiparameter graphs of LmrP 9C-404C, 188C-404C, 98C-374C, 164C-312C and 164C-374C measured at pH 8, pH 5, pH 8+Hoechst (H) and pH 8 + roxithromycin (R).**

Each panel represents a 2D plot with a 1D histogram on each axis. **A** Donor lifetime, **B** Acceptor lifetime and **C** stoichiometry, versus *E*. **D** Donor lifetime versus Donor anisotropy and **E** Acceptor lifetime versus Acceptor anisotropy. The red line in **A** is the static FRET line calculated as described in the Methods, and the red lines in **D** and **E** are the lines calculated via the Perrin equations. The correction factors and the parameters are given in **F**.

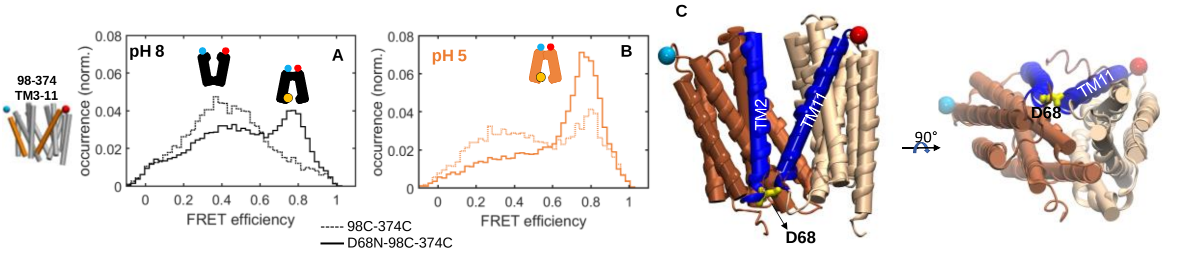


**Fig S.3: Modulation of the conformational equilibrium at the extracellular side by the D68N mutation.**

The *E* histograms measured at (**A**) pH8 and (**B**) pH5 are shown in black and orange respectively, without (dashed line) or with the D68N mutation (solid line). **C:** Localisation of the D68 residue on the protein. The N- and C-halves are coloured in dark and light brown respectively, and the probes are represented in blue/red. The D68 residue is situated at the bottom of the protein in yellow. A closer look at this region shows that this residue interacts with TM11. The effect of this mutation on the transport activity of the distance reporter 98C-374C is given in Fig. S1F.

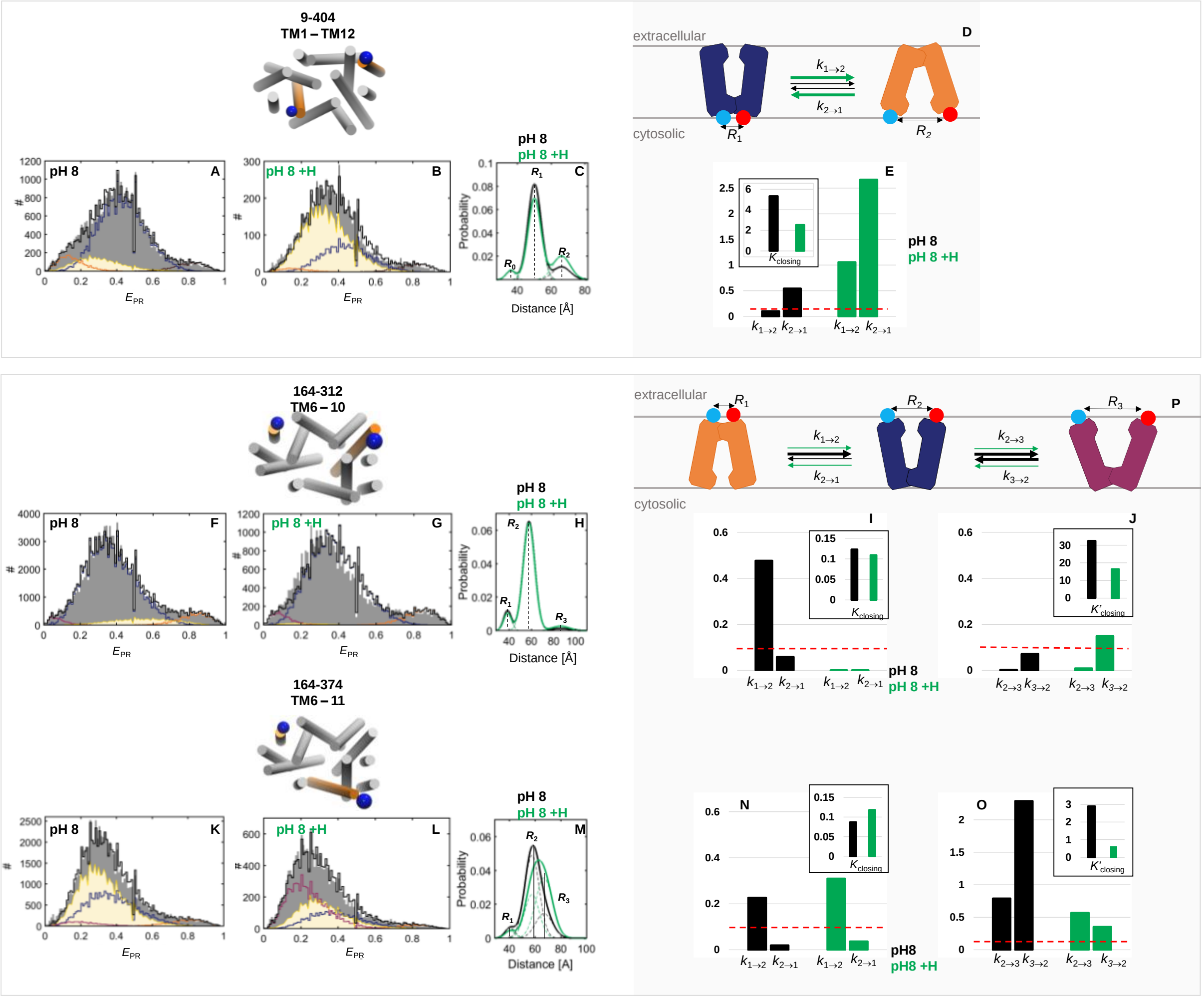


**Fig S.4: Effect of Hoechst binding on the transitions**

**A-B**: Experimental $E_{PR}$ histograms (gray bars) of LmrP 9C-404C burst data rebinned in 1-ms bins and acquired at pH 8 in (**A**) absence or (**B**) presence of Hoechst. The total PDA model (black stairs) is a sum of the inward-open-state bins (orange stairs), inward-closed-state bins (blue stairs), inward-very-closed-state bins (brown stairs) and bins describing state-interconverting molecules (yellow highlighted area). **C**: The probability density plots of the PDA data from panels A (black solid line) and B (green solid line) are a sum of the underlying components (dashed lines), and are characterized by the distances $R_0$, $R_1$ and $R_2$. **D.** Cartoon depicting the conformational interconversion kinetics scheme at the cytosolic side (blue = inward-closed, orange = inward-open). **E**: Bar charts of the opening ($k_{1\rightarrow2}$) and closing ($k_{2\rightarrow1}$) rate constants (in $ms^{-1}$) (black = apo, green = Hoechst). The dashed red line is the rate constant below which interconversion kinetics cannot be determined accurately. The inset represents the equilibrium constant for closing.

**F-G**: Experimental $E_{PR}$ histograms (gray bars) of LmrP 164C-312C burst data rebinned in 1-ms bins and acquired at pH 8 in (**F**) absence or (**G**) presence of Hoechst. The total PDA model (black stairs) is a sum of the outward-closed-state bins (orange stairs), outward-open-state bins (blue stairs), extra-open-state bins (purple stairs) and bins describing state-interconverting molecules (yellow highlighted area). **H**: The probability density plots of the PDA data from panels A (black solid line) and B (green solid line) are a sum of the underlying components (dashed lines), and are characterized by the distances $R_1$, $R_2$ and $R_3$. **I**: Bar charts of the opening ($k_{1\rightarrow2}$) and closing ($k_{2\rightarrow1}$) rate constants (in $ms^{-1}$) (black = apo, green = Hoechst). **J**: Bar charts of the extra-opening ($k_{2\rightarrow3}$) and closing ($k_{3\rightarrow2}$) rate constants (in $ms^{-1}$) (black = apo, green = Hoechst). The dashed red lines are the rate constant below which interconversion kinetics cannot be determined accurately. The insets represent the equilibrium constant for closing.

**K-L**: Experimental $E_{PR}$ histograms (gray bars) of LmrP 164C-374C burst data rebinned in 1-ms bins and acquired at pH 8 in (**K**) absence or (**L**) presence of Hoechst. The total PDA model (black stairs) is a sum of the outward-closed-state bins (orange stairs), outward-open-state bins (blue stairs), extra-open-state bins (purple stairs) and bins describing state-interconverting molecules (yellow highlighted area). **M**: The probability density plots of the PDA data from panels A (black solid line) and B (green solid line) are a sum of the underlying components (dashed lines), and are characterized by the distances $R_1$, $R_2$ and $R_3$. **N**: Bar charts of the opening ($k_{1\rightarrow2}$) and closing ($k_{2\rightarrow1}$) rate constants (in $ms^{-1}$) (black = apo, green = Hoechst). **O**: Bar charts of the extra-opening ($k_{2\rightarrow3}$) and closing ($k_{3\rightarrow2}$) rate constants (in $ms^{-1}$) (black = apo, green = Hoechst). The dashed red lines are the rate constant below which interconversion kinetics cannot be determined accurately. The insets represent the equilibrium constant for closing. **P.** Cartoon depicting the conformational interconversion kinetics scheme at the cytosolic side (orange = outward-closed, blue = outward-open, purple = outward-extra-open).

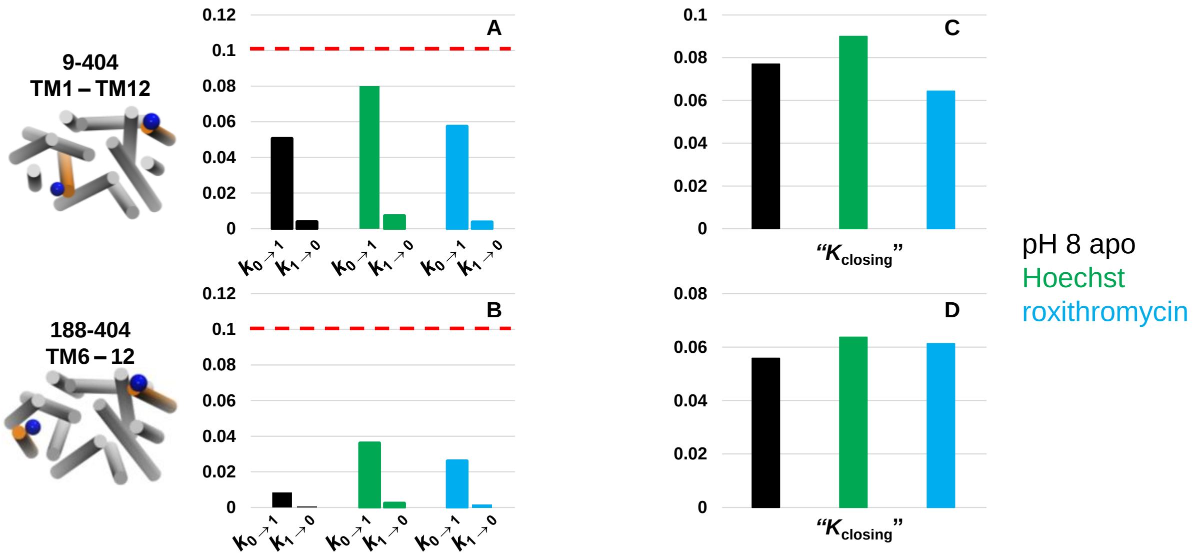


**Fig S.5: No interconversion detected between the high FRET state and the inward-closed state**
Left: the two constructs probing the cytosolic side, bottom view. **A-B:** bar charts of the constant rates (in $ms^{-1}$) from the inward-very-closed to the inward-closed state ($k_{0\text{-}1}$) and from the inward-closed to the inward-very-closed state ($k_{1\text{-}0}$) on the cytosolic side, for LmrP 9C-404C and 188C-404C respectively, pH 8 in black, pH 8 + Hoechst in green and pH 8 +roxithromycin in blue. The dashed red lines are the rate constant below which interconversion kinetics cannot be determined accurately. **C-D:** ratio between the rate constants $k_{1\text{-}0}$ and $k_{0\text{-}1}$ , for LmrP 9C-404C and 188C-404C respectively.

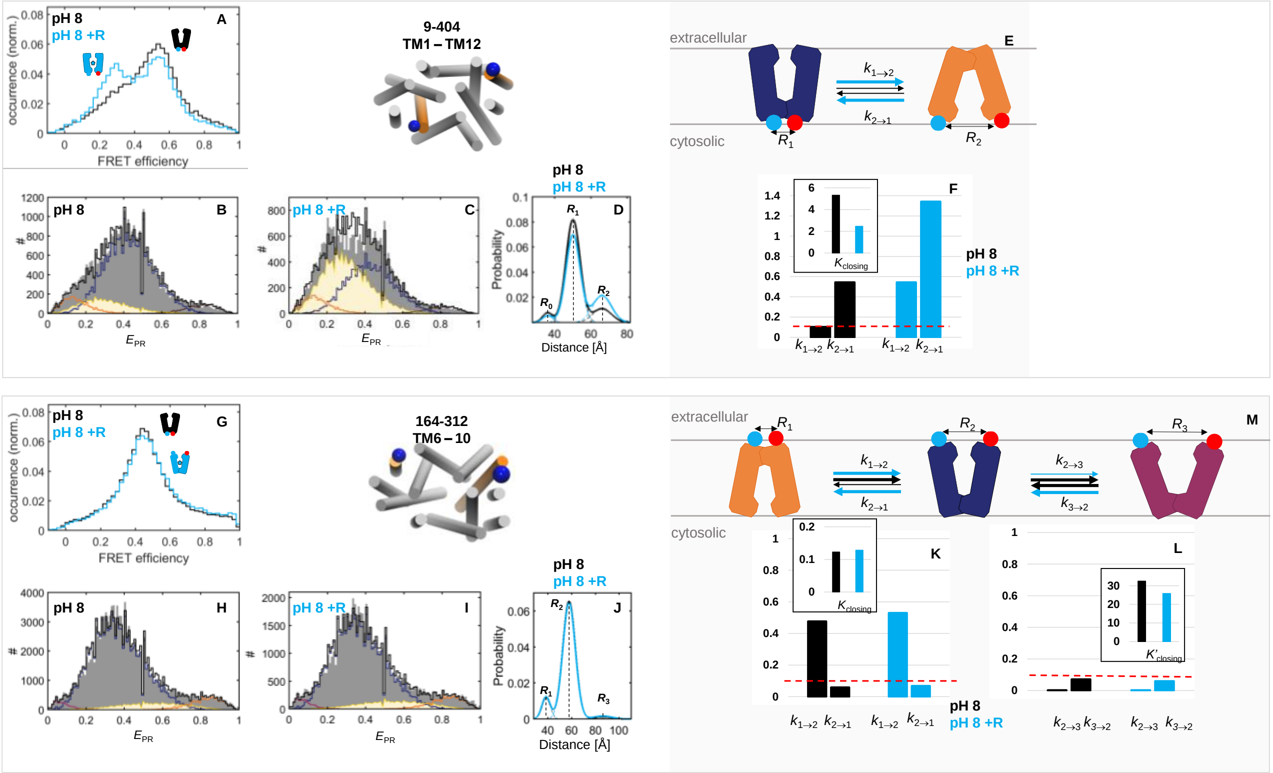


**Fig S.6: Effect of roxithromycin binding on the transitions.**

**A:** *E* histograms at pH 8 (black) and with roxithromycin (light blue) for LmrP 9C-404C. **B-C**: Experimental $E_{PR}$ histograms (gray bars) of LmrP 9C-404C burst data rebinned in 1-ms bins and acquired at pH 8 in (**B**) absence or (**C**) presence of roxithromycin. The total PDA model (black stairs) is a sum of the inward-open-state bins (orange stairs), inward-closed-state bins (blue stairs), inward-very-closed-state bins (brown stairs) and bins describing state-interconverting molecules (yellow highlighted area). **D**: The probability density plots of the PDA data from panels A (black solid line) and B (light blue solid line) are a sum of the underlying components (dashed lines), and are characterized by the distances $R_0$, $R_1$ and $R_2$. **E.** Cartoon depicting the conformational interconversion kinetics scheme at the cytosolic side (blue = inward-closed, orange = inward-open). **F**: Bar charts of the opening ($k_{1\rightarrow2}$) and closing ($k_{2\rightarrow1}$) rate constants (in $ms^{-1}$) (black = apo, light blue = roxithromycin). The dashed red line is the rate constant below which interconversion kinetics cannot be determined accurately. The inset represents the equilibrium constant for closing.

**G:** *E* histograms at pH 8 (black) and with roxithromycin (light blue) for LmrP 164C-312C. **H-I**: Experimental $E_{PR}$ histograms (gray bars) of LmrP 98C-374C burst data rebinned in 1-ms bins and acquired at pH 8 in (**H**) absence or (**I**) presence of roxithromycin. The total PDA model (black stairs) is a sum of the outward-closed-state bins (orange stairs), outward-open-state bins (blue stairs), extra-open-state bins (purple stairs) and bins describing state-interconverting molecules (yellow highlighted area). **J**: The probability density plots of the PDA data from panels A (black solid line) and B (light blue solid line) are a sum of the underlying components (dashed lines), and are characterized by the distances $R_1$, $R_2$ and $R_3$. **K**: Bar charts of the opening ($k_{1\rightarrow2}$) and closing ($k_{2\rightarrow1}$) rate constants (in $ms^{-1}$) (black = apo, light blue = roxithromycin). **L**: Bar charts of the extra-opening ($k_{2\rightarrow3}$) and closing ($k_{3\rightarrow2}$) rate constants (in $ms^{-1}$) for all extracellular reporter mutants (black = apo, light blue = roxithromycin). The dashed red line is the rate constant below which interconversion kinetics cannot be determined accurately. The insets represent the equilibrium constant for closing. **M.** Cartoon depicting the conformational interconversion kinetics scheme at the cytosolic side (orange = outward-closed, blue = outward-open, purple = outward-extra-open).

**Supp. Table 1: The PDA results of the two cytosolic distance reporters measured at pH 8, with Hoechst or with roxithromycin**

| | 9C-404C | | | 188C-404C | | |
|---|---|---|---|---|---|---|
| | **pH 8** | **Hoechst** | **Roxi.** | **pH 8** | **Hoechst** | **Roxi.** |
| $R_0$ **(Å)** | 36.7 | 36.7 | 36.7 | 35.1 | 35.1 | 35.1 |
| $R_1$ **(Å)** | 50.3 | 50.3 | 50.3 | 48.2 | 48.2 | 48.2 |
| $R_2$ **(Å)** | 65.8 | 65.8 | 65.8 | 63.9 | 63.9 | 63.9 |
| $k_{0\rightarrow1}$ **(ms$^{-1}$)** | 0.051* | 0.08* | 0.057* | 0.008* | 0.036* | 0.026* |
| $k_{1\rightarrow0}$ **(ms$^{-1}$)** | 0.004* | 0.007* | 0.004* | 0.000* | 0.002* | 0.002* |
| **“$K_{closing}$” $k_{1\rightarrow0}$ / $k_{0\rightarrow1}$** | 0.077 | 0.09 | 0.064 | 0.056 | 0.064 | 0.061 |
| **Dwell time$_0$ (ms)** | 19.8* | 12.5* | 17.4* | 119.0* | 27.7* | 38.3* |
| **Dwell time$_1$ (ms)** | 256.4* | 138.9* | 270.3* | >>>* | 434.8* | 625.0* |
| $k_{1\rightarrow2}$ **(ms$^{-1}$)** | 0.102 | 1.059 | 0.544 | 0.005* | 0.411 | 0.731 |
| $k_{2\rightarrow1}$ **(ms$^{-1}$)** | 0.543 | 2.667 | 1.342 | 0.009* | 0.644 | 0.865 |
| $K_{closing}$ | 5.322 | 2.518 | 2.468 | 2.022 | 1.566 | 1.183 |
| **Dwell time$_1$ (ms)** | 9.8 | 0.9 | 1.8 | 217.4* | 2.4 | 1.4 |
| **Dwell time$_2$ (ms)** | 1.8 | 0.4 | 0.7 | 107.5* | 1.5 | 1.1 |
| **$\chi^2$ (1ms time bin)** | 4.59 | 2.72 | 8.37 | 4.18 | 2.91 | 8.30 |

*The value lies outside the sensitive range of values that can be determined via burst analysis.

With $R_0$ corresponding to the inter-dyes distance of the ‘inward-very-closed’ state, $R_1$ of the inward-closed state and $R_2$ of the inward-open state. The dwell times in each of these three states is inferred from the rate constants, with $k_{0\rightarrow1}$ being the opening rate constant from the $R_0$ state to the $R_1$ state, $k_{1\rightarrow0}$ being the closing rate constant from the $R_1$ state to the $R_0$ state, $k_{1\rightarrow2}$ being the opening rate constant from the $R_1$ state to the $R_2$ state and $k_{2\rightarrow1}$ being the closing rate constant from the $R_2$ state to the $R_1$ state. $K_{closing}$ is the equilibrium constant for closing as defined in the Methods. For each mutant, the four time windows (0.2, 0.5, 0.75 and 1ms) and the three distances were globally optimized between the pH 8 apo, + Hoechst and + roxithromycin data.

**Supp. Table 2: The PDA results of the three extracellular distance reporters measured at pH 8, with Hoechst or with roxithromycin**

| | 98C-374C | | | 164C-312C | | | 164C-374C | |
|---|---|---|---|---|---|---|---|---|
| | **pH 8** | **Hoechst** | **Roxi.** | **pH 8** | **Hoechst** | **Roxi.** | **pH 8** | **Hoechst** |
| **$R_1$ (Å)** | 39.9 | 43.5 | 41.9 | 38.5 | 38.5 | 38.5 | 40.6 | 40.6 |
| **$R_2$ (Å)** | 56.8 | 59.2 | 57.3 | 57.9 | 57.9 | 57.9 | 57.6 | 57.6 |
| **$R_3$ (Å)** | 76.5 | 74.9 | 71.6 | 86.4 | 86.4 | 86.4 | 67.0 | 67.0 |
| **$k_{1\rightarrow2}$ ($ms^{-1}$)** | 0.483 | 0.006* | 0.963 | 0.477 | 0.000* | 0.530 | 0.227 | 0.310 |
| **$k_{2\rightarrow1}$ ($ms^{-1}$)** | 0.079* | 0.001* | 0.246 | 0.058* | 0.000* | 0.068* | 0.020* | 0.037* |
| **$K_{closing}$** | 0.190 | 0.158 | 0.235 | 0.123 | 0.109 | 0.128 | 0.087 | 0.118 |
| **Dwell $time_1$ (ms)** | 2.1 | 169.5* | 1.0 | 2.1 | >>>* | 1.9 | 4.4 | 3.2 |
| **Dwell $time_2$ (ms)** | 12.6* | 769.5* | 4.1 | 17.1* | >>>* | 14.7* | 50.8* | 27.3* |
| **$k_{2\rightarrow3}$ ($ms^{-1}$)** | 0.250 | 0.054* | 0.053* | 0.002* | 0.009* | 0.002* | 0.791 | 0.569 |
| **$k_{3\rightarrow2}$ ($ms^{-1}$)** | 1.023 | 0.223 | 0.212 | 0.071* | 0.149 | 0.059* | 2.288 | 0.354 |
| **$K'_{closing}$** | 3.084 | 2.482 | 3.697 | 32.454 | 16.406 | 25.869 | 2.894 | 0.623 |
| **Dwell $time_2$ (ms)** | 4.0 | 18.4* | 18.8* | 454.5* | 109.9* | 434.8* | 1.3 | 1.7 |
| **Dwell $time_3$ (ms)** | 1.0 | 4.3 | 4.7 | 14.0* | 6.7 | 16.8* | 0.4 | 2.8 |
| **$\chi^2$ (1ms time bin)** | 3.65 | 6.01 | 14.65 | 18.95 | 15.61 | 12.66 | 19.4 | 2.93 |

*The value lies outside the sensitive range of values that can be determined via burst analysis.

With $R_1$ corresponding to the inter-dyes distance of the outward-closed state, $R_2$ of the outward-open state and $R_3$ of the outward-extra-open state. The dwell times in each of these three states is inferred from the rate constants, with $k_{1\rightarrow2}$ being the opening rate constant from the $R_1$ state to the $R_2$ state, $k_{2\rightarrow1}$ being the closing rate constant from the $R_2$ state to the $R_1$ state, $k_{2\rightarrow3}$ being the opening rate constant from the $R_2$ state to the $R_3$ state and $k_{3\rightarrow2}$ being the closing rate constant from the $R_3$ state to the $R_2$ state. $K_{closing}$ and $K'_{closing}$ are the equilibrium constant for closing as defined in the Methods. For each mutant, the four time windows (0.2, 0.5, 0.75 and 1ms) were globally optimized between pH 8 apo, + Hoechst and + roxithromycin data.
